# A tug of war between filament treadmilling and myosin induced contractility generates actin ring

**DOI:** 10.1101/2021.06.06.447254

**Authors:** Qin Ni, Kaustubh Wagh, Aashli Pathni, Haoran Ni, Vishavdeep Vashisht, Arpita Upadhyaya, Garegin A. Papoian

## Abstract

In most eukaryotic cells, actin filaments assemble into a shell-like actin cortex under the plasma membrane, controlling cellular morphology, mechanics, and signaling. The actin cortex is highly polymorphic, adopting diverse forms such as the ring-like structures found in podosomes, axonal rings, and immune synapses. The biophysical principles that underlie the formation of actin rings and cortices remain unknown. Using a molecular simulation platform, called MEDYAN, we discovered that varying the filament treadmilling rate and myosin concentration induces a finite size phase transition in actomyosin network structures. We found that actomyosin networks condense into clusters at low treadmilling rates or high myosin concentration but form ring-like or cortex-like structures at high treadmilling rates and low myosin concentration. This mechanism is supported by our corroborating experiments on live T cells, which exhibit ring-like actin networks upon activation by stimulatory antibody. Upon disruption of filament treadmilling or enhancement of myosin activity, the pre-existing actin rings are disrupted into actin clusters or collapse towards the network center respectively. Our analyses suggest that the ring-like actin structure is a preferred state of low mechanical energy, which is, importantly, only reachable at sufficiently high treadmilling rates.

## 1 Introduction

In eukaryotic cells, actin filaments and myosin motors self-organize into a diversity of shapes (*1*). A shell-like cortex is ubiquitously found under the cell membrane, which is characterized by a mesh-like geometry and plays an indispensable role in defining cellular shape and mechanochemical responses (*2–4*). In immune cells such as T cells, the actin cortex reorganizes into a peripheral quasi-2D actin ring that sequesters different signaling complexes in separate concentric domains upon stimulation by antigen-presenting cells (*5–9*). Ring-like actin geometries have also been widely found in other sub-cellular structures such as podosomes and axons (*10, 11*). How actin filaments and associated motors and proteins assemble into such ubiquitous networks that enable the shape control of living cells and tissues remains poorly understood due to the complexity and non-equilibrium nature of actomyosin networks.

*In vitro* networks reconstituted from purified proteins have been extensively used to derive the minimal set of determining conditions that govern the assembly, growth and structural properties of actin networks. However, *in vitro* networks exhibit strikingly different higher order structures compared to cellular networks. In notable contrast to the ring-like or shell-like networks ubiquitously seen in living cells, *in vitro* experiments primarily result in actomyosin networks comprised of clusters that originate from global geometric collapse due to myosin motor driven contractility (*12–19*). The origins of this disparity likely lies in the qualitatively different parameter spaces occupied by *in vitro* actin networks compared to cellular networks.

Actin filaments are highly dynamic, undergoing rapid polymerization and depolymerization, and are subject to contractile forces generated by myosin motors (*20,21*). Actin polymerization is polarized: monomeric actin (G-actin) binds to the barbed ends of filaments and polymeric actin (F-actin) dissociates from the pointed ends in a process called treadmilling (*22–24*). We hypothesized the differences between the predominant actomyosin architectures formed *in vitro* versus those observed *in vivo* may arise from the large difference in the corresponding treadmilling rates: *in vitro* networks reconstituted from purified proteins exhibit treadmilling rates that are often several-fold slower than those observed *in vivo* due to the lack of regulators that promote actin filament polymerization and disassembly (*25–29*). We further postulated that these differences in treadmilling rates render *in vitro* networks less resistant to myosin-induced collapse. A systematic way to explore how treadmilling rates and myosin contractility combine to shape actomyosin network architecture is essential to probe our hypothesis. This is a difficult experimental task, requiring careful manipulation of molecular machinery and actin polymerization kinetics. Such limitations can be overcome by computer simulations, which provide a powerful way to capture the complex chemistry and mechanics of the active cytoskeleton, and bring significant mechanistic insights.

In order to find a minimal set of conditions that lead to the formation of rings and cortices, we combined computer simulations via the open-access platform MEDYAN (Mechanochemical Dynamics of Active Networks) (*15*) and experiments on live T cells. We find that the competition between actin filament treadmilling and myosin contractility determines the overall network morphology. Our simulations showed that the speed of actin filament treadmilling drives the network away from global centripetal actomyosin clustering, resulting instead in centrifugal condensation that creates ring-like and cortex-like structures, without tethering filaments to the boundary. On the other hand, increasing myosin motor activity or decreasing filament treadmilling rates lead to centripetal collapse of actin networks, creating clusters in the network center. Our corroborating experiments on live T cells and simulations mimicking experimental conditions showed that, indeed, hyper-activating myosin II via Calyculin A (CalyA) or inhibiting filament treadmilling via Latrunculin A (LatA) disassembled pre-existing actin rings, causing the network to condense centripetally, resulting in clusters.

Furthermore, our computational analysis indicates that actin filaments located at the network periphery have lower mechanical energy as compared to those that form actomyosin clusters and hence represent the energetically preferred configuration. However, this energetic state is only achievable at sufficiently high treadmilling rates, while at lower treadmilling rates, the system gets trapped in long-lived metastable states where actin filaments instead condense into clusters. In summary, our work shows that a tug of war between filament treadmilling and myosin-induced contraction determines the fate of actomyosin architectures: the energetically favorable ring/cortex states are kinetically accessible only at higher treadmilling rates. Our findings reveal that the assembly and stability of various cellular actin structures are crucially regulated by the fine-tuning of filament treadmilling, which can be achieved by the activation of accessory proteins, such as formin, profilin, and cofilin, via local biochemical signaling.

## 2 Results

### 2.1 Dissecting and modeling the T cell actin ring

In order to construct a molecular model of actin rings, we first examined the F-actin distribution in live Jurkat T cells expressing TD-Tomato-F-tractin (an indirect reporter of F-actin) and EGFP-MLC (myosin light chain). These cells were allowed to spread on an activating glass surfaces coated with anti-CD3 antibody and imaged with time-lapse total internal reflection fluorescence (TIRF) microscopy to visualize the dynamics of actin reorganization (Video 1). Upon activation by stimulatory antibodies, the actin cytoskeleton in T cells reorganizes into a ring-like structure characterizing the immune synapse (Fig. 1a-c and Ref. 5–9). The actin ring consists of an outer lamellipodial region and an inner lamellar ring. In the outer ring, Arp2/3 is activated by WASP near the membrane (*30*), generating a branched actin network that largely excludes NMII (Fig. 1c). The inner ring is enriched in actin filaments decorated with NMII which form actomyosin “arcs” (Fig. 1b-c). The central region is largely depleted in actin and NMII.

**Figure 1:**
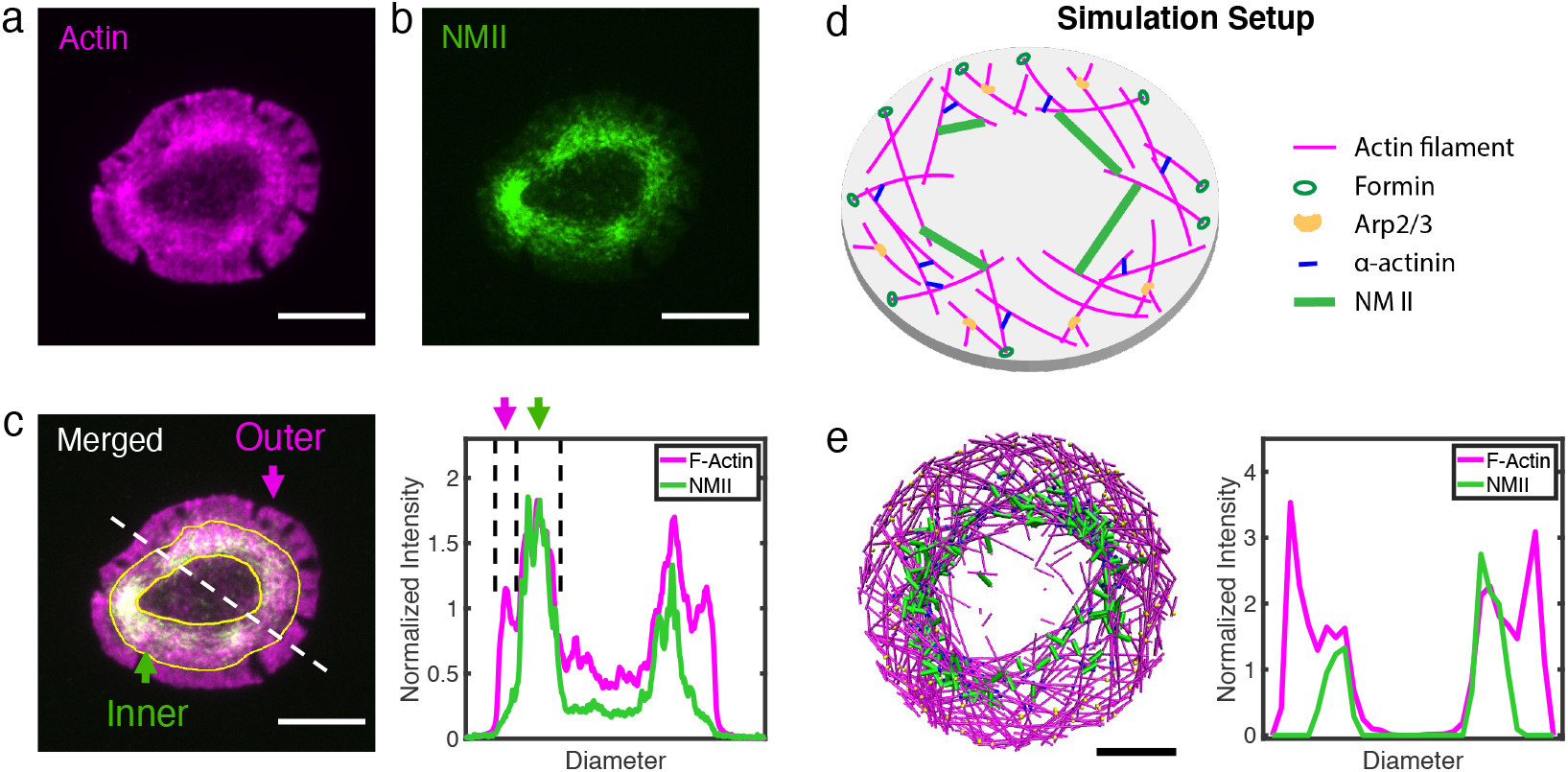
Actin and NMIIs distribution of actin rings in T cells and simulation. (a-b) Representative snapshots of actin (a) and NMII (b) in actin rings of live Jurkat T cells on acti-CD3 antibody. Actin is labeled by tdTomato F-tractin (magenta), and NMII is labeled by MLC-EGFP (green). (c) The distribution of actin and NMII within the T cell actin ring. (a-b) Scale bar = 5 µm. (d) Setup of simulations using MEDYAN with major cytoskeletal components labeled. (e) A representative snapshot of simulation and the corresponding distribution of actin and NMII along the diametral direction. *C*_*actin*_ = 120*µM, C*_*NMII*_ = 0.1*µM, C*_*alpha*−*actinin*_ = 4*µM, C*_*Arp*2*/*3_ = 1*µM, andC*_*formin*_ = 0.3*µM*. Scale bar = 1 µm.

To understand the biophysical determinants of ring formation and stability, we modeled the formation of actin ring systems using MEDYAN, a simulation platform that combines sophisticated, single molecule level treatment of cytoskeletal reactions, polymer mechanics, and mechanochemical feedback. Actin networks were simulated in a thin oblate cylinder with diameters between 3.8 µm and 10 µm, to mimic the lateral dimensions of small mammalian cells (Fig. 1d). Model details can be found in Methods and Supplementary Text 1. We first modeled an actin network with Arp2/3 mediated branching near the periphery. Simulations show that this preferential activation of branching alone is sufficient to generate a lamellipodia-like actin ring, similar to the outer T cell ring, without any other cytoskeletal components or filament tethering to the boundary (Fig. S1). We then added the motor protein NMII, crosslinker alpha-actinin, and a filament nucleator formin, which are essential components for actin network remodeling and are ubiquitously found in actin rings and cortices (*1, 2*). To mimic the *in vivo* conditions, we excluded NMIIs from the peripheral region with Arp2/3 mediated branching. Upon tuning the concentrations of cytoskeletal components and filament treadmilling rates, we found that the network self-organizes into and maintains an outer lammellipodia-like ring and an inner lamellar-like ring with similar actomyosin spatial distribution as the actin ring in T cells (Fig. 1e). Also similar to T cells (*5–7*), simulated F-actin undergoes retrograde flow due to filament polymerization against the boundary and NMII generated contraction (Fig. S2). Even without spatial restrictions on actin or myosin at the periphery, our simulated networks resemble the inner actomyosin ring found in T cells, suggesting that the formation of ring-like actin structure is a consequence of actomyosin self-organization. We next focused on the origins of this inner actomyosin ring.

### 2.2 Building a minimal model for actin ring formation

To explore the minimal determinants for actin ring formation, we first modeled networks with only actin filaments at different average treadmilling rates (⟨*r*_*TM*_⟩ = 0.57 s^-1^, 1.41 s^-1^, and 2.21 s^-1^) based on actin filament assembly kinetics reported from prior experiments (*25, 31, 32*).These systems also include formin at a concentration of 100 nM (*24, 33*). We found that disordered actin networks were created at all treadmilling conditions tested (Fig. S3b). We quantified the spatiotemporal evolution of the network geometry by plotting the median of the radial filament density distribution (*R*_*median*_) as a function of time (Fig. S3a). In NMII-free networks, we observed a relatively uniform filament density across the network regardless of treadmilling rates (Fig. S3c). In this case, the network geometry is dominated by stochastic filament treadmilling that is not spatially biased. The boundary plays an important role, as the boundary repulsion force inhibits barbed end polymerization such that filaments reaching the boundary rapidly depolymerize and eventually disassemble. The loss of filaments through depolymerization is compensated by the nucleation of new filaments, resulting in dynamic and disordered structures (Video 2).

We next explored how these disordered networks behaved upon the introduction of crosslinking and motor contractility. We allowed the network to evolve for 300 s at different ⟨*r*_*TM*_⟩ as described above to reach a steady disordered state, and then added NMIIs and the actin crosslinker alpha-actinin to generate contractile forces. The addition of NMIIs and crosslinkers changed the steady state network geometry, as measured by *R*_*median*_ (Fig. 2a). For slow treadmilling rates (⟨*r*_*TM*_⟩ = 0.56*s*^−1^), the addition of NMIIs and alpha-actinin resulted in the clustering of actin filaments (Fig. 2b-iii, and Video 2). This geometric pattern is consistent with prior *in vitro* and *in silico* studies on contractile actomyosin networks (*12–14,16*), where contractility can be defined as a symmetry breaking event accompanied by a geometric collapse of the network. The average local concentration of actin within the clusters was 234 µM, which is almost six-fold higher than the initial G-actin concentration (40 µM), suggesting a high degree of condensation. Although the size and location of actin clusters varied significantly across multiple trajectories (Fig. S4), a decreasing *R*_*median*_ suggests that the overall collapse is centripetal (Fig. 2b-iii).

**Figure 2:**
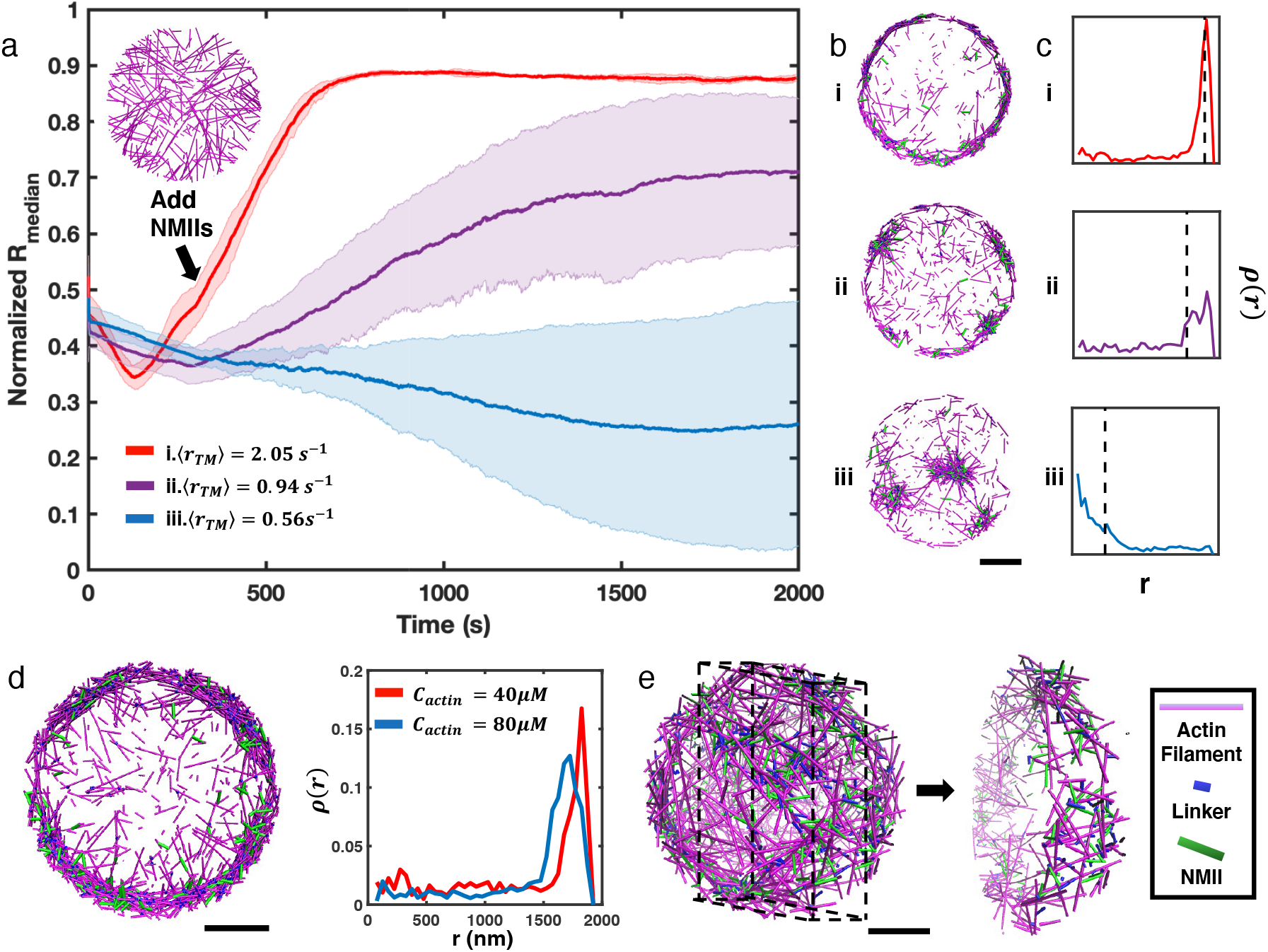
NMII contractility induces geometric collapse of treadmilling actin filaments. (a) Normalized medians of radial filament density distribution (*R*_*median*_) at different tread-milling rates (⟨*r*_*TM*_⟩) are shown. The treadmilling rate is defined as the average number of actin monomers added per filament per second at the barbed ends - equivalent to the rate of F-actin depletion from the pointed ends - after reaching the kinetic steady state (Supplementary Text 2). 0.06 µM of NMII and 4 µM of alpha-actinin were added at 301 s. The inset figure is a snapshot at t = 300 s of networks with ⟨*r*_*TM*_⟩= 2.05*s*^−1^. The shaded error bars represent the standard deviation across 5 runs. (b-c) Representative snapshots at each treadmilling condition (b) and their radial filament density distribution, *ρ*(*r*) (c) are shown. Dashed lines in (c) indicate the position of *R*_*median*_. (d) Representative snapshot of ring-like networks with 80 µM actin (left), and *ρ*(*r*) of actin rings with 40 µM actin and 80 µM actin are shown (⟨*r*_*TM*_⟩= 1.35 s^-1^). (e) A snapshot of a spherical cortex-like network (left) and a slice showing the internal structure (right). (a,b,d,e) Actin filaments are magenta cylinders, NMIIs are green cylinders and linkers are blue cylinders in all snapshots. All scale bars are 1 µm.

Our simulations suggest that actin networks are subject to two competing processes: treadmilling, which tends to homogeneously distribute filaments in the network, and NMII mediated contractility, which tends to trap filaments into clusters. We thus explored changes in the actin network geometry by increasing the treadmilling rate while maintaining the same concentration of NMII. Although filament nucleation occurs stochastically throughout the entire network and there is no filament tethering near the boundary, we discovered that after the addition of NMII to rapidly treadmilling networks (⟨*r*_*TM*_⟩ = 2.05*s*^−1^), filaments steadily accumulate at the network boundary (Fig. 2a-i, and Video 2). During this process, we observed that NMIIs deformed many filaments and gradually changed their orientation from being perpendicular to the boundary to parallel (Video 3). Upon allowing the system to further evolve for several hundred seconds, we found that actin networks transformed into ring-like structures (Fig. 2b-i). Networks with intermediate ⟨*r*_*TM*_⟩ = 0.94*s*^−1^ form a mixture of clusters and rings (Fig. 2b-ii, and Video 2).

The resulting actin rings are highly condensed, with a thickness of a few hundred nanometers and exhibiting local actin concentrations similar to those found in actin clusters (263 µM, Supplementary Text 3). Increasing the initial G-actin concentration increases the thickness of actin rings (Fig. 2d). Most filaments in actin rings are oriented parallel to the boundary (Fig. S5), forming small actin clusters that undergo azimuthal flow (Video 2-3). Analogous ring-like patterns were observed on a larger system with a diameter of 10 µm (Fig. S6). In a spherical system, networks evolved into hollow spherical cortex-like geometries under similar conditions (Fig. 2e, Fig. S7, and Video 4).

### 2.3 Competition between filament treadmilling and NMII contractility determines network morphology

To further examine how treadmilling rate regulates the formation of distinct actomyosin architectures, we performed extensive simulations at different treadmilling rates. Indeed, ⟨*r*_*TM*_⟩ emerges as a key control parameter that governs the steady state network geometry. Below a critical ⟨*r*_*TM*_⟩, which is 0.94 *s*^−1^ in our simulations, networks geometrically collapse into clusters, while above this critical ⟨*r*_*TM*_⟩, they preferentially evolve into ring-like geometries (Fig. 3a and b). The radial distribution of the ring state is characterized by higher *R*_*median*_ and smaller standard deviation compared with the cluster phase. Interestingly, *R*_*median*_ as a function of ⟨*r*_*TM*_⟩ displays a sharp increase as the network transitions from the cluster state to the ring state (Fig. 3b). Because the *R*_*median*_ trajectories after adding NMIIs are almost linear before reaching a steady state, we quantified the network remodeling speed by measuring the slopes of the linear part of the *R*_*median*_ trajectories. We found that the network remodeling speed is positively correlated with ⟨*r*_*TM*_⟩ (Fig. 3c), indicating that ⟨*r*_*TM*_⟩ is an important factor driving network structural evolution.

**Figure 3:**
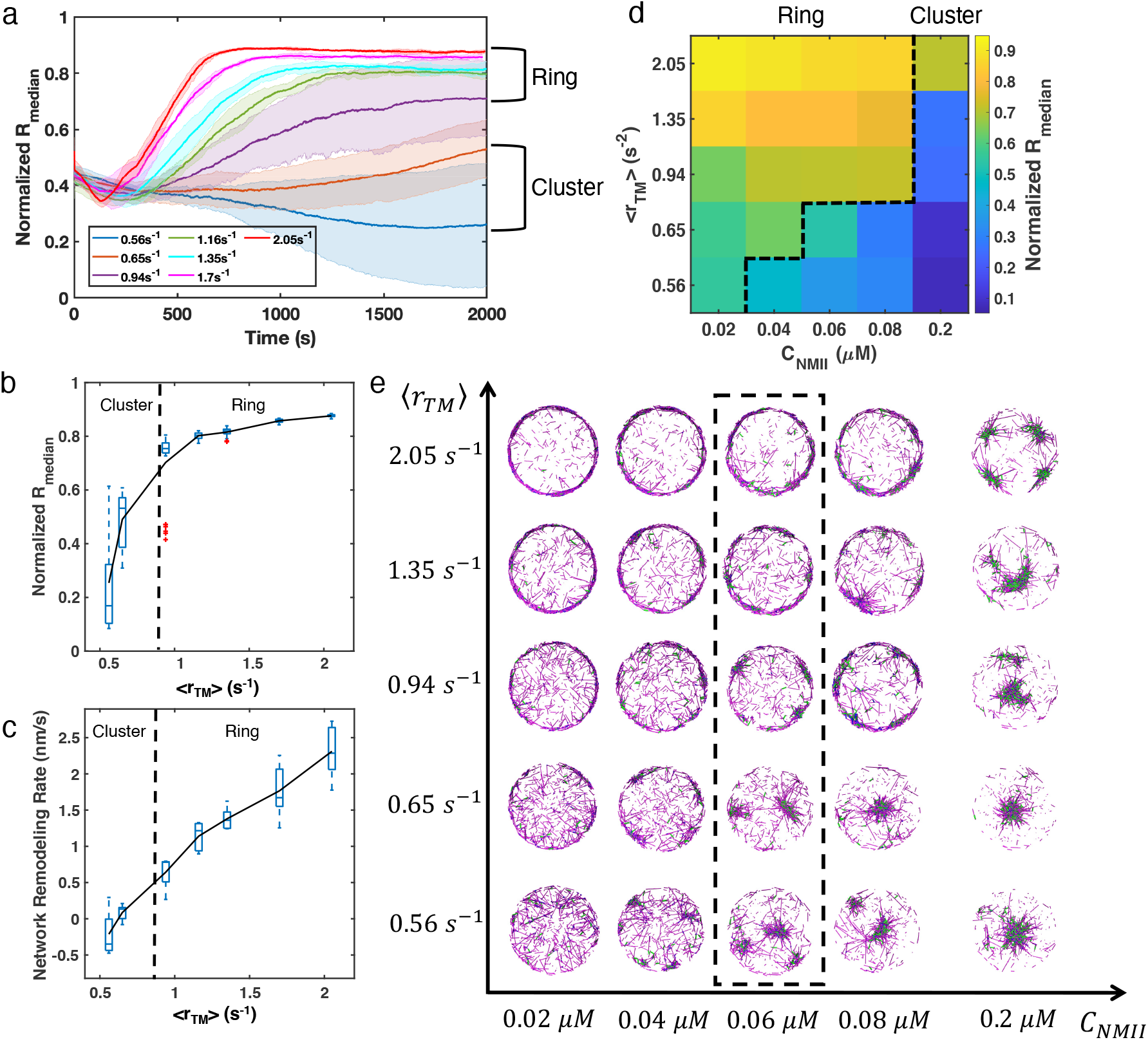
Treadmilling rates and NMII concentrations regulated network structural transition. (a) Normalized medians of radial filament density distribution (*R*_*median*_) as a function of time at different ⟨*r*_*TM*_⟩ (0.56 *s*^−1^ to 2.05 *s*^−1^) are shown. The shaded colors represent the standard deviation of means for 5 runs. (b) The box plot shows the average *R*_*median*_ at the last 500 seconds of simulation at each treadmilling rate. Solid line connects the mean *R*_*median*_ at each ⟨*r*_*TM*_⟩. (c)The box plot shows the speed of network remodeling, measured as the slope of linear part of *R*_*median*_ after 300 seconds. The solid line connects the mean remodeling rates at each ⟨*r*_*TM*_⟩. (a-c) *C*_*actin*_ = 40*µM, C*_*NMII*_ = 0.06*µM, C*_*alpha*−*actinin*_ = 4*µM*. (d) Steady state *R*_*median*_ at different ⟨*r*_*TM*_⟩ (0.56 *s*^−1^ to 2.05 *s*^−1^) and *C*_*NMII*_ (0.02 to 0.2 µM). (e) Representative snapshots of steady state actin network structures at different ⟨*r*_*TM*_⟩ and *C*_*NMII*_. Representative snapshots of trajectories in (a) are shown in the dash box. (d-e) *C*_*actin*_ = 40*µM, C_alpha−actinin_* = 4*µM*.

We next varied the NMII concentration (*C*_*NMII*_) at different treadmilling rates, obtaining a phase diagram delineating actin network morphologies (Fig. 3d-e). The phase diagram indicates that the critical ⟨*r*_*TM*_⟩ for actin ring formation increases as *C*_*NMII*_ increases. Networks collapse into clusters for ⟨*r*_*TM*_⟩ below the critical value. The higher *C*_*NMII*_ is, the more likely a cluster tends to localize to the geometric center of the network, indicating that NMII induced contractility drives the centripetal condensation. When ⟨*r*_*TM*_⟩ and *C*_*NMII*_ are both low, the network becomes disordered (for example, see Fig. 3e, ⟨*r*_*TM*_⟩ = 0.56*s*^−1^ and *C*_*N*_ *MII* = 0.02*µM*). Similarly, increasing network contractility by tuning alpha-actinin concentration also results in a transition from ring-like networks at low linker concentrations to bundles and clusters at higher concentrations (Fig. S8).

### 2.4 Inhibition of actin dynamics disrupts actin rings in live cells and *in silico*

In order to further understand how treadmilling regulates ring-like actin networks, we experimentally disrupted F-actin dynamics in live Jurkat T cells expressing EGFP-F-tractin. Since it is not feasible to directly control the treadmilling rate in experiments, we used the actin inhibitor, Latrunculin-A (LatA), which decreases the polymerization rate and increases the depolymerization rate by sequestering G-actin and accelerating phosphate release from ADP-Pi-actin (*32, 34, 35*). Upon the formation of the actin ring at the contact zone, LatA (at different concentrations) was added to spreading cells and the resulting effect on the rings was monitored with time-lapse imaging. In order to compare with simulations, we used the fluorescence intensity as a reporter of F-actin levels and calculated a normalized *R*_*median*_ to quantify the evolution of the actin network under varying degrees of LatA inhibition compared to vehicle control (Fig. 4a). With weak inhibition (*C*_*LatA*_ =250 nM), the ring like structure is perturbed but largely preserved for several minutes (Fig. 4b, and Video 5). At higher doses of LatA (*C*_*LatA*_ =500 nM and 1 µM, Fig. 4c, and Video 5), *R*_*median*_ rapidly decreases (Fig. 4d), indicating a collapse of the network towards the geometric center of the cell. The rate of centripetal collapse of the actin network increases with increasing *C*_*LatA*_ (Fig. 4e). The dismantling of the actin ring is also accompanied by the formation of F-actin clusters or bundles (Fig. 4b-c).

**Figure 4:**
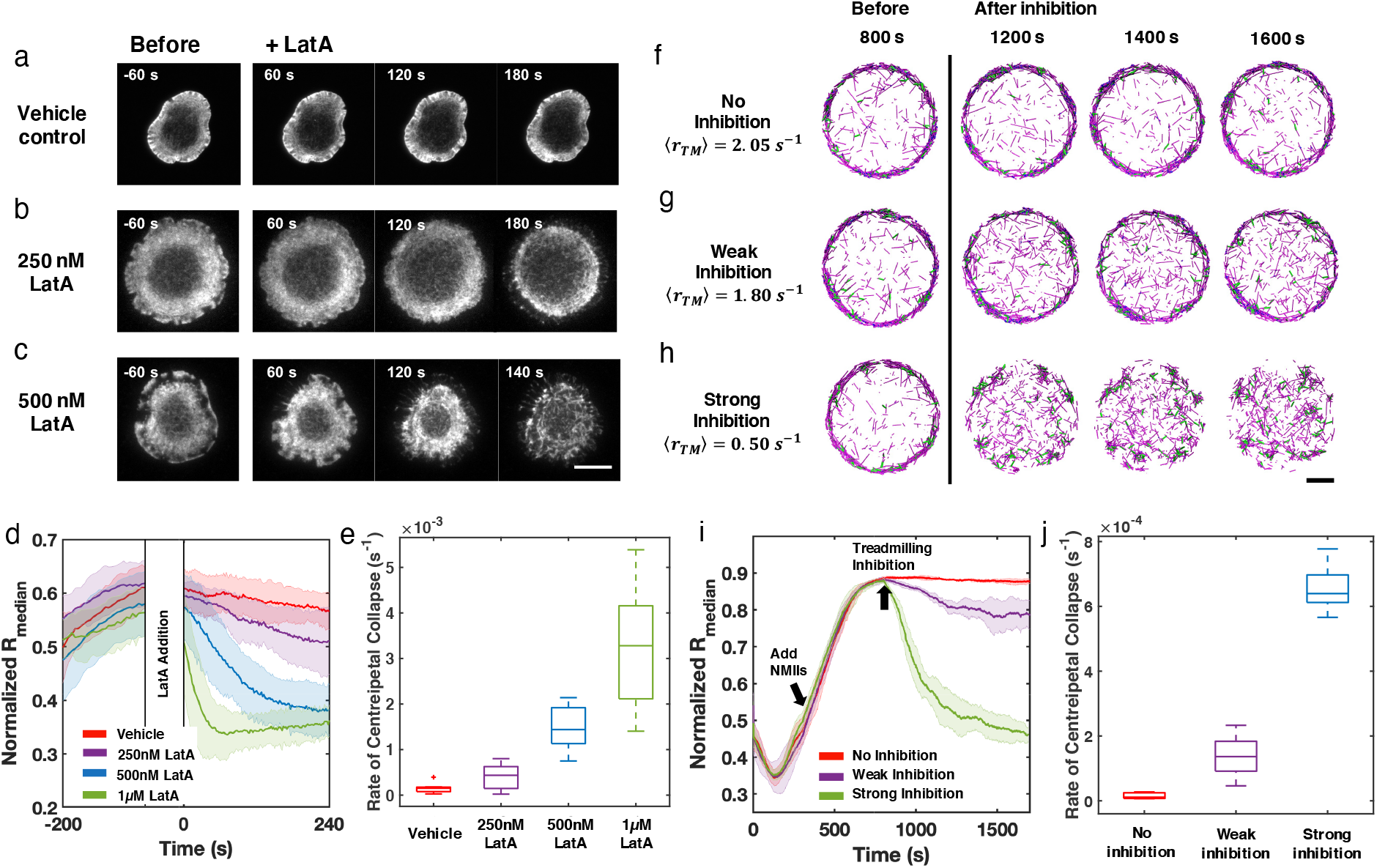
Inhibition of actin dynamics induces collapse of actin rings in live T cells and *in silico*. (a-c) Timelapse montages of Jurkat T cells expressing F-tractin-EGFP spreading on anti-CD3 coated glass substrates. Cells were treated with (a) vehicle control (0.1% DMSO), (b) 250 nM LatA, or (d) 500 nM LatA between 300 and 360 seconds after contact with activating surface. The first post-treatment image is labeled as 0s. Timelapse images illustrate the centripetal collapse of the actin ring upon treatment with LatA. Timescales of this collapse depend on the concentration of LatA as can be seen from the timestamps on the images. Scale bar is 10 µm. (d) Quantification of the spatial organization of the actin network using the normalized median of radial filament density distribution. Shaded error bars represent the standard deviations across trajectories (7-11 cells per condition). (e) Box plots showing the rate of centripetal collapse, measured as the slope of the *R*_*median*_ distribution after inhibition. (f-h) Timelapse montages of simulations of (f) control, (g) weak inhibition, and (h) strong inhibition. Treadmilling rate in these conditions are 2.05*s*^−1^, 1.80*s*^−1^, and 0.50*s*^−1^, respectively. Indicated ⟨*r*_*TM*_⟩ is the averaged treadmilling from 1500s to the end of simulations. Scale bar indicates 1 µm. (i) Medians of radial filament density distribution at different conditions. (j) Rate of centripetal collapse rate, measured as the slope of the *R*_*median*_ distribution after inhibition. (i, j) The shaded color and error bars represent the standard deviation across trajectories, n = 5 runs per condition.

To compare with these experimental observations, we perturbed actin network assembly *in silico* after ring-like networks are established. Based on recent work on reconstituted actin networks under LatA treatment (*32*), we reduced the polymerization rate constants and increased the depolymerization rate constants to mimic the effect of LatA on G-actin sequestering and accelerating depolymerization to closely follow the T cell experiments (see Supplementary Text 4 for model details). Actin rings (no inhibition, Fig. 4f) were created in the same way as shown in Figure 2a-i. Upon the formation of stable actin rings at 800 seconds, we perturbed actin filament polymerization to different extents to mimic weak and strong LatA inhibition (Video 6). We found that actin rings persist under weak inhibition (Fig. 4g), while they collapse into clusters under strong inhibition (Fig. 4h). The disruption of actin filament assembly also reduces ⟨*r*_*TM*_⟩ from 2.05*s*^−1^ to 1.80*s*^−1^ (weak inhibition) and 0.50*s*^−1^ (strong inhibition), respectively. Measurements of *R*_*median*_ and the rate of collapse (Fig. 4i and j) at different inhibition conditions reveal the centripetal collapse of the ring network, reproducing the above-described experimental observations.

### 2.5 Enhancement of NMII activity leads to centripetal contraction of actomyosin rings in T cells and *in silico*

In order to validate the role of NMII activity in regulating ring-like actin networks, we next altered the NMII dynamics in live Jurkat T cells. Under vehicle control (DMSO), actin rings are relatively stable over the timescale of 10 minutes, and the F-actin distribution displays a steep transition from a depletion zone at the cell center to a high-intensity plateau (Fig. S9a). Calyculin A (CalyA) application to enhance NMII activity (*36*) leads to an increase in contractility and a centripetal collapse of the actin network (Fig. 5a, and Video 7), as quantified by the decrease of *R*_*median*_ Fig. 5c. On the other hand, upon treatment with Y-27632, an inhibitor of NMII upstream regulator Rho kinase (*37*) which decreases myosin based contractility, the network becomes more disordered and displays a shallower transition from the central depletion zone to the peripheral plateau (Fig. 5b, and Video 7). We quantified these changes by calculating the slope of the normalized F-actin intensity from the center to plateau region (Supplementary Text 5). As shown in Fig. 5d, he slope remained constant over time under vehicle application, while it decreased upon Y-27632 application, indicating that the network becomes more diffusive and disordered, and the ring integrity is compromised with loss of myosin contractility. These results confirm that NMII is a central regulator of actin network structure, and high NMII activity is antagonistic to actin ring formation.

**Figure 5:**
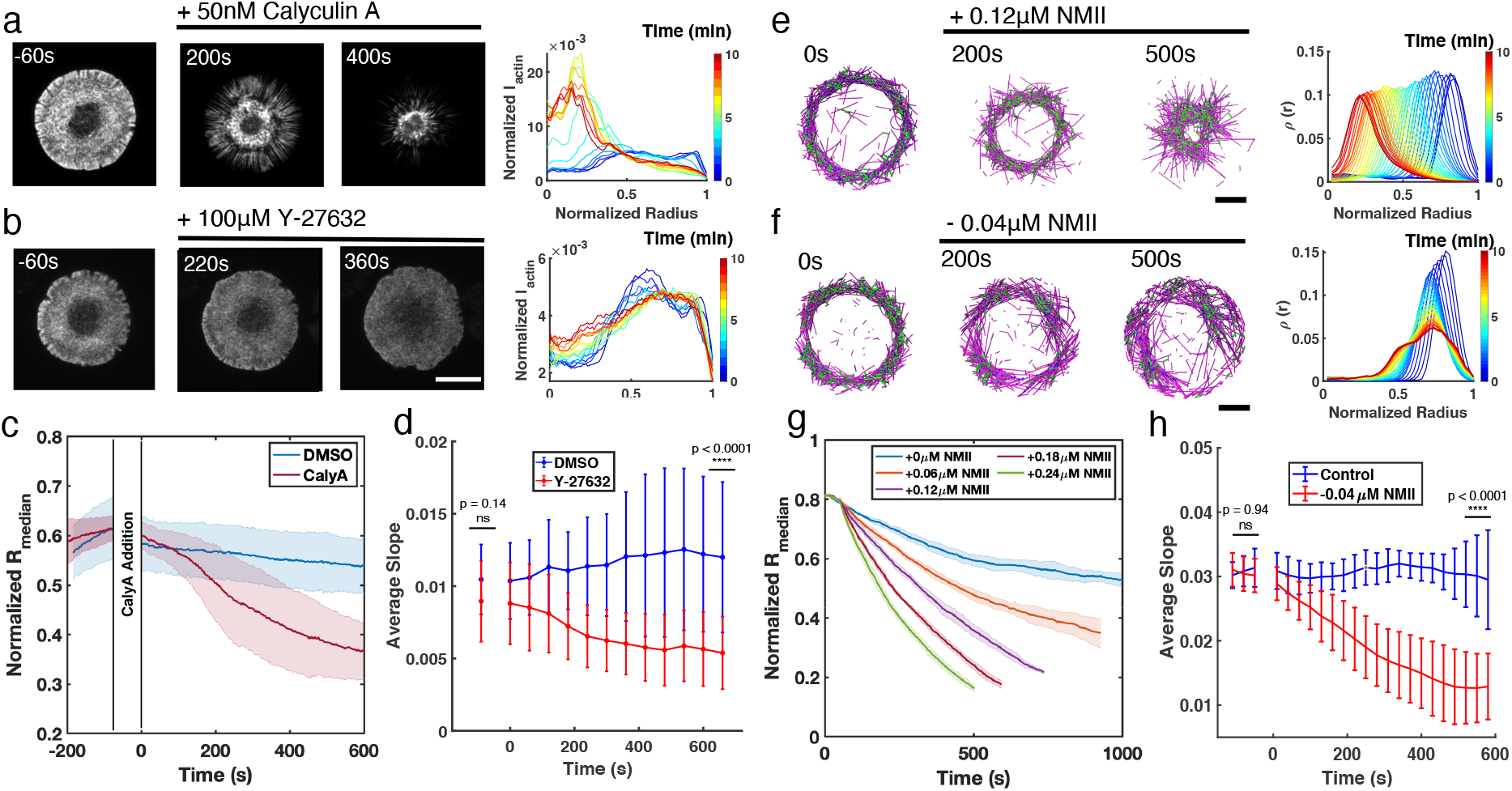
Enhancement or inhibition of NMII regulates actin structure in live T cells and *in silico*. (a-b) Time lapse montages of Jurkat T cells expressing F-tractin-EFGP spreading on anti-CD3 coated glass substrates (left) and the normalized radial filament density distribution *ρ*(*r*) (right). After achieving maximal spreading, cells were treated with (a) 50 nM CalyA, and (b) 100 µM Y-27632. Scale bar is 10 µm. (c) The normalized median of radial filament density distribution *R*_*median*_. n = 12 for vehicle (0.5% DMSO), and 14 for CalyA. Two sample t-test was performed for the first point before drug addition (ns, p=0.83) and 600 s after drug addition (****,p¡0.0001).(d) The slope of the intensity profiles over the transition region from the center to the peripheral plateau as a function of time. n= 25 for vehicle (0.1% DMSO), and 24 for Y-27832. Two sample t-test was performed for the first point (before drug addition) and the last point (660 s after drug addition). (e-f) Timelapse montages of simulations(left) and *ρ*(*r*) at different times (right) mimicking actin rings in (e) CalyA treatment by increasing NMII levels, and (f) Y-27632 treatment by reducing NMII levels. The control condition is shown in Figure. S9, containing 80 µM actin, 0.18 µM NMII, and 4 µM alpha-actinin. Scale bar is 1 µm. (g) The evolution of *R*_*median*_ for different levels of NMII addition. Blue curve is control, and other curves are simulations with indicated levels of NMII added to mimic the CalyA experiment. (h) The slope of the intensity profiles over the transition region from the center to the peripheral plateau as a function of time for simulations of Y-27632 addition. Blue curve is control while orange curve represents simulations after reduction of NMII concentration by 0.04 µM. Two sample t-test was performed for the first three points (before inhibition) and the last three points (510 s to 600 s after drug addition). (g-h) n = 5 runs per condition. (a-h) In all figures, 0s represented the first time point recorded after drug addition (for experiments) or NMII addition/depletion (for simulations). (c,d,g,h) Shaded color and error bar represent the standard deviation across cells or simulation trajectories.

We then validated the role of NMII in shaping actin structure using MEDYAN simulations. To reduce the computational time, we first initialized actin ring networks and then increased or decreased NMII concentrations. Under control conditions, we tuned the actin and NMII concentrations to mimic the evolution of the actin ring in T cells (Fig. S9b). In remarkable agreement with experiments, enhancing NMII levels induces centripetal collapse of the network (Fig. 5e, and Video 8), and the speed of the collapse is proportional to the amount of NMII added to the system (Fig. 5f). These results also indicate that a confined boundary is not required for the maintenance of actin rings. On the other hand, upon reduction of NMII levels, the actin ring becomes more disordered (Video 8) and the slope of the center to plateau F-actin distribution decreases (Fig. 5g and h), in agreement with Y-27632 inhibition experiments.

### 2.6 Energetic origins of structural polymorphism in active networks

We next explored the chemical and mechanical properties of actin networks at various treadmilling rates. We found that the numbers of F-actin filaments, bound linkers, and bound motors remain nearly constant across different ⟨*r*_*TM*_⟩, while distributions of diffusive molecules, such as G-actin and nucleators, also did not show spatial localization, being uniformly distributed throughout the simulation volume (Fig. S10). These observations suggest that ring-like architectures do not form because of the enrichment of soluble constituent molecules near the periphery.

The lack of enrichment of soluble molecules in the periphery suggested a possible energetic origin of the structures. We thus examined the mechanical energy (*U*_*Mech*_) of the system, which primarily arises from filament bending in our simulation. For fixed concentrations of NMII (0.06 µM) and crosslinker (4 µM), we found that *U*_*Mech*_ decreases with increasing ⟨*r*_*TM*_⟩ (Fig. 6a). In addition, *U*_*Mech*_ undergoes a sharp reduction when ⟨*r*_*TM*_⟩ reaches the critical threshold, with *U*_*Mech*_ of actin rings being 2-3 fold lower than that of clusters. Moreover, we found that *U*_*Mech*_ is negatively correlated with *R*_*median*_, regardless of the structural state (Fig. 6b). Since higher *R*_*median*_ indicates localization of actin filaments at the network periphery, this negative correlation indicates that configurations with the lowest mechanical energy are those with a ring-like geometry. These results suggest that the peripheral arrangement of actin filaments is more energetically favorable than more distorted configurations found in centripetal clusters.

**Figure 6:**
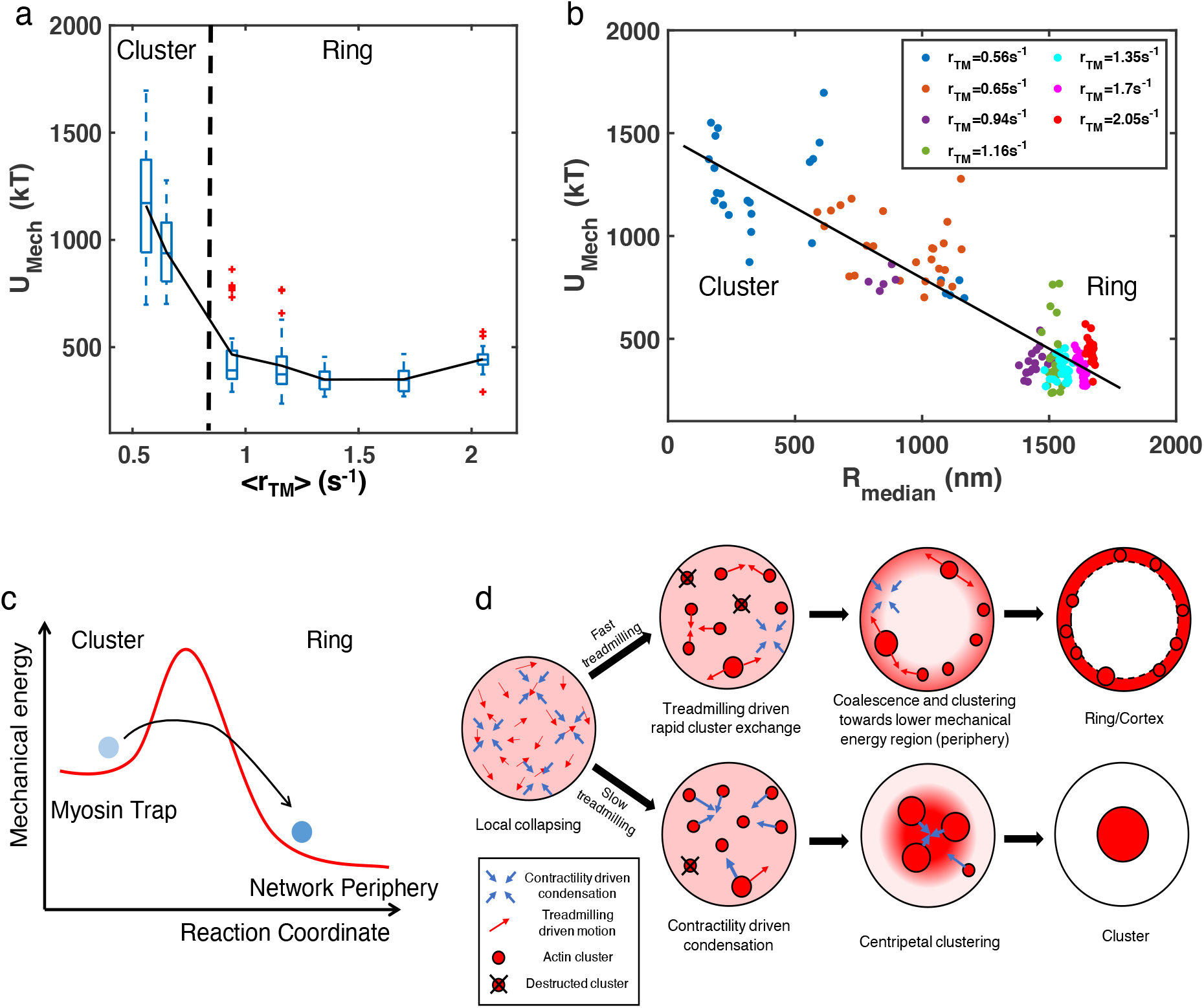
Energetic origins of actin rings. (a) The box plot shows the steady state *U*_*Mech*_ at each treadmilling rate. *U*_*Mech*_ is the sum of the bending energy of actin filaments and the stretching energy of filaments, motors, and linkers. Solid line connects the mean *U*_*Mech*_ at each ⟨*r*_*TM*_⟩. (b) Mechanical energy (*U*_*Mech*_) and the corresponding *R*_*median*_ at different *r*_*TM*_. Each data point represents the average *U*_*Mech*_ and *R*_*median*_ per 100 seconds of the last 500 seconds of simulation. (a-b) *C*_*actin*_ = 40*µM, C*_*NMII*_ = 0.06*µM, C*_*alpha*−*actinin*_ = 4*µM*, with varying ⟨*r*_*TM*_⟩as shown in Figure 3a-c. n = 5 runs per condition. (c) A graphical description showing the envisioned energy landscape for generating actin cortices. (d) Schematic showing the formation of actin ring/cortex *versus* clusters. At low treadmilling rates, networks are dominated by myosin-driven contraction, leading to centripetal collapse into clusters (lower). Faster filament treadmilling allows networks to overcome the myosin-driven centripetal motion, where filaments tend to move to the network periphery due to lower energy (upper).

## 3 Discussion

Detailed mechanochemical modeling using MEDYAN shows that active actin networks exhibit a striking morphological transition upon changes in the filament treadmilling rate. We found that two distinct types of dynamic structures emerge due to the interplay between treadmilling rates and NMII contractility in an initially disordered network: (1) actin clusters formed in slow-treadmilling or high *C*_*NMII*_ networks and (2) ring-like and cortex-like structures spontaneously assembled in fast-treadmilling and low *C*_*NMII*_ networks. This geometric transition does not require filament tethering to the boundary or a spatially biased filament assembly. We also observed a sharp transition in the system’s mechanical energy during the transformation from a multi-cluster network to a ring architecture. Such a sharp change in morphology and mechanical energy, induced by tuning filament treadmilling speed, is indicative of a finite size phase transition.

While phase transitions in many biomolecular systems are often driven by passive biomolecular interactions (*38, 39*), in this work we identified a phase transition in cytoskeletal networks that is induced by non-equilibrium actomyosin dynamics. Our analysis shows that the formation of actin rings and cortices arises from the competition between filament treadmilling and myosin induced contraction. The addition of myosin motors and crosslinkers to an initially disordered actin network induces contractile forces, creating energetically metastable actomyosin clusters (Fig. 6c). Rapid filament treadmilling provides a mechanism for escaping these traps (*15, 27, 40*), giving rise to smaller clusters that rapidly dissolve and reappear. In this state, the network has more freedom to remodel its structure in order to lower the mechanical energy. Indeed, our analysis suggests that as the actin filament distribution shifts to the network periphery, the smaller curvature at the boundary results in a decrease in filament bending, thereby lowering the mechanical energy of the network (Fig. 6a-b). As a consequence, actin filaments at high treadmilling speeds rapidly accumulate at the network periphery, contributing to the build up of an actin ring in flattened volumes or actin cortices in fully 3D spherical geometries (Fig. 6d, upper). In contrast, networks undergoing slow filament treadmilling are trapped in cluster-like configurations that have higher mechanical energy. The latter networks are dominated by myosin-driven contractility, leading to a highly non-ergodic state in which actin filaments undergo centripetal collapse (Fig. 6d, lower).

Although some previous computational studies have studied how network morphology and contractility are regulated by treadmilling rates (*17, 40, 41*), the active formation of ring-like or cortical shell-like networks and their underlying mechanisms have not been explained before. In this work, we have shown that rapid treadmilling and the presence of myosin are sufficient to create ring-like or cortex-like actomyosin networks in a system with confined boundaries. Furthermore, it is likely that filament binding to the cell membrane (*42*) or the spatially biased localization of actin assembly regulators, such as Arp2/3 (*9*), can further enhance the formation of ring-like structures. Studying these and other regulatory processes will bring new mechanistic insights into the organization and dynamics of cortices/rings and their defects, which occur in primary immunodeficiencies, autoimmune disorders, and cancers.

## Methods

### Simulation setup

In this work, we employed an open-access mechanochemical platform for simulating active matter (MEDYAN (*15*)) to investigate the spatiotemporal evolution of actin networks under different treadmilling and myosin motor conditions. MEDYAN accounts for two overlapping phases and their interactions. 1) Diffusing G-actin and unbound formins, NMII and linkers are spatially dissolved in a solution phase. In this phase, the network is discretized into compartments based on the Kuramoto length of G-actin, which is the mean-free path that G-actin molecules are expected to diffuse before undergoing their next reaction (*43*). Diffusing chemical species are assumed to be well-mixed within each compartment, and inter-compartment transports are modeled as stochastic diffusion reactions. 2) Polymeric filaments and bound species comprise the continuous polymeric phase which is overlaid on the solution phase. The polymeric phase is mechanically active, where filament bending, stretching, and steric interactions are taken into account. Bound motors and linkers are modeled as harmonic springs based on the mechanical properties of NMII and alpha-actinin. A boundary repulsion potential restricts filaments within the volume boundary. Filament polymerization is affected by interactions with the boundary, following the Brownian Ratchet model (*44*). The following chemical reactions stochastically occur among the two phases: filaments can polymerize, depolymerize, and interact with myosin and crosslinker; formins are able to bind to G-actin and nucleate filaments; filaments that are only two monomers long can be rapidly destroyed. The chemical reaction modeling engine is based on an efficient and statistically accurate Next Reaction Method (NRM) (*45*), which is a variant of the Gillespie Algorithm (*46*).

We initialized *de novo* cytoskeletal networks in MEDYAN with small seed filaments, 40 µM diffusing G-actin, and 100 nM filament nucleators based on their reported cytoplasmic concentrations (*47*). Most of the simulations were carried out in a thin oblate geometry, having a diameter ranging from 3.8 µm to 10 µm and an effective height of 200 nm. The spherical simulation volume has a diameter of 4 µm. We tuned the barbed end polymerization rate and pointed end depolymerization rate to model the effects of treadmilling promoters such as formin, profilin, and cofilin. To monitor the actual speed of treadmilling, we define ⟨*r*_*TM*_⟩ as the average barbed end elongation rate, which is also equal to the shortening rate of the pointed end at steady state. Networks were allowed to assemble with only filament polymerization, depolymerization, nucleation, and disassembly for 300 seconds. At 300 s, 0.06 µM NMII and 4 µM alpha-actinin crosslinkers are added. The local density of clusters and rings were measured using a customized density based clustering algorithm. Further simulation details can be found in Supplementary Text 1.

### Cell culture and transfection

E6.1 Jurkat T cells were grown in RPMI medium supplemented with 10% Fetal Bovine Serum (FBS) and 1% penicillin-streptomycin at 37°C in a CO_2_ incubator. Transfections were performed with 2 × 10^5^ cells using 1 µg of plasmid by electroporation using a Neon electroporation kit (Thermo Fisher Scientific). Prior to imaging, cells were transferred to CO_2_ independent L-15 medium (Fisher Scientific).

### Plasmids and reagents

pEGFP-C1 F-tractin-EGFP was a gift from Dyche Mullins (Addgene plasmid # 58473) (*48*). The td-Tomato-F-tractin plasmid was a gift from Dr. John A. Hammer and the MLC-EGFP plasmid was a gift from Dr. Robert Fischer, National Heart, Lung, and Blood Institute. Latrunculin A was purchased from Sigma Aldrich and its solvent, Calyculin A was purchased from Cell Signaling Technology, Y-27632 was purchased from Selleck Chemicals, and dimethyl sulfoxide (DMSO) was purchased from Thermo Fisher Scientific.

### Preparation of glass coverslips

Sterile 8-well chambers (Cellvis) were incubated with 0.01% poly-L-lysine solution in distilled water for 10 minutes and then dried at 37°C for 1 hour. Poly-L-lysine coated chambers were then incubated with anti-human CD3 antibody (HIT3a clone, Thermo Fisher Scientific) in PBS at a concentration of 10 µg/mL for 2 hours at 37°C or overnight at 4°C. Following incubation, the chambers were washed 5 times with L-15 and warmed prior to imaging. For inhibitor experiments with Calcyulin-A and Y-27632, Jurkat T cells transfected with F-Tractin-EGFP were seeded and allowed to activate on anti-CD3 coated glass coverslips. 50 nM Calyculin-A, 100 uM Y-27632 or vehicle (DMSO) were added after the cells had formed an actin ring. Fluorescence images were acquired as above with a frame interval of 2 seconds using a 100X, 1.49 NA oil immersion objective.

### Microscopy

Transfected T cells were seeded on anti-CD3 coated glass coverslips and allowed to activate for 5 minutes. Chambers were maintained at 37°C using a stage-top incubator (Okolab). Latrunculin A or vehicle (DMSO) were added at specified concentrations 5 minutes after seeding the cells. Fluorescence and interference reflection microscopy (IRM) images were acquired using an inverted microscope (Ti-E, Nikon, Melville, NY) with a scientific CMOS camera (Prime BSI, Photometrics, Tucson, AZ) with a frame interval of 2 seconds. F-tractin-EGFP was imaged using total internal reflection fluorescence (TIRF), using a 60X, 1.49 NA oil immersion objective. One background image was captured during every session in order to perform background subtraction.

### Image analysis

Initial preprocessing of images was done using Fiji (*49*). A custom MATLAB script was written to perform background subtraction. The IRM or actin images were used to find the outline and centroid of the cells. 50 uniformly spaced lines were drawn from the centroid and these 50 line profiles were pooled together to generate a histogram of intensities as a function of a normalized distance to the centroid. The median of the distribution of intensities (and hence F-actin) was estimated for each time point.

## Acknowledgments

We would like to thank A. Chandrasekaran, C. Floyd, and J. Komianos for helpful discussions and feedback on the manuscript. This work was supported by National Science Foundation grants CHE-1800418 and PHY-1806903. A.U. acknowledges support from the grants PHY 1607645 and NIH R01 GM131054. Computational resources were provided by Deepthought2 HPC at University of Maryland.

## Supplementary Figures

**Figure S1:**
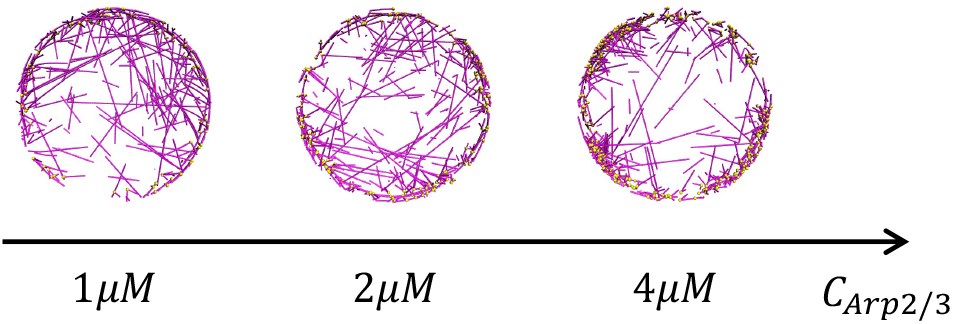
Representative snapshots of actin networks that consists of 40 µM actin and 1-4 µM Arp2/3 without crosslinkers or motors. Arp2/3 (represented as yellow beads) is activated 500 nm away from the boundary.

**Figure S2:**
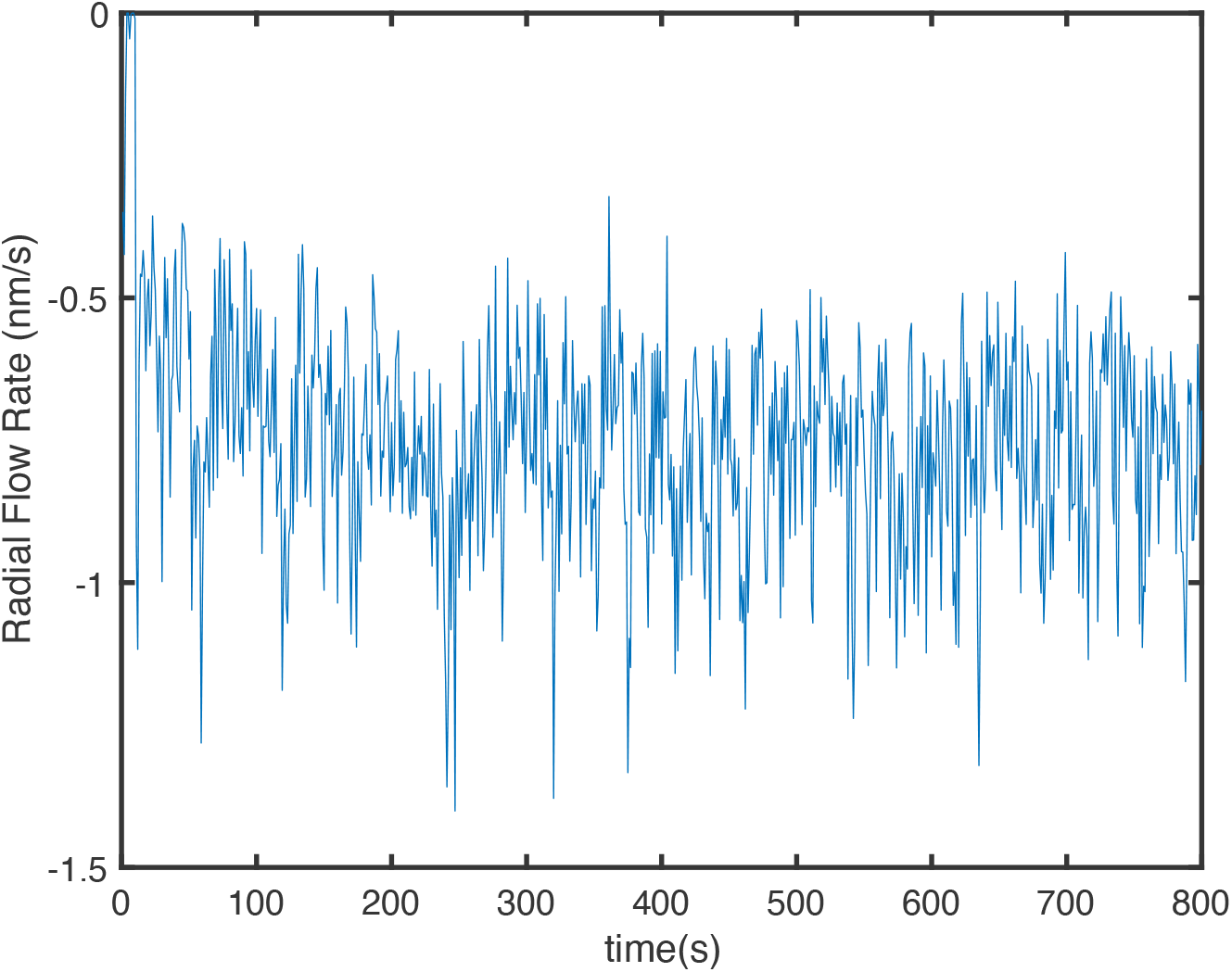
F-actin flow rate along the radial direction. Flow rate is quantified by tracking the mean displacement of individual F-actin molecules. Negative flow rate indicates the direction is centripetal.

**Figure S3:**
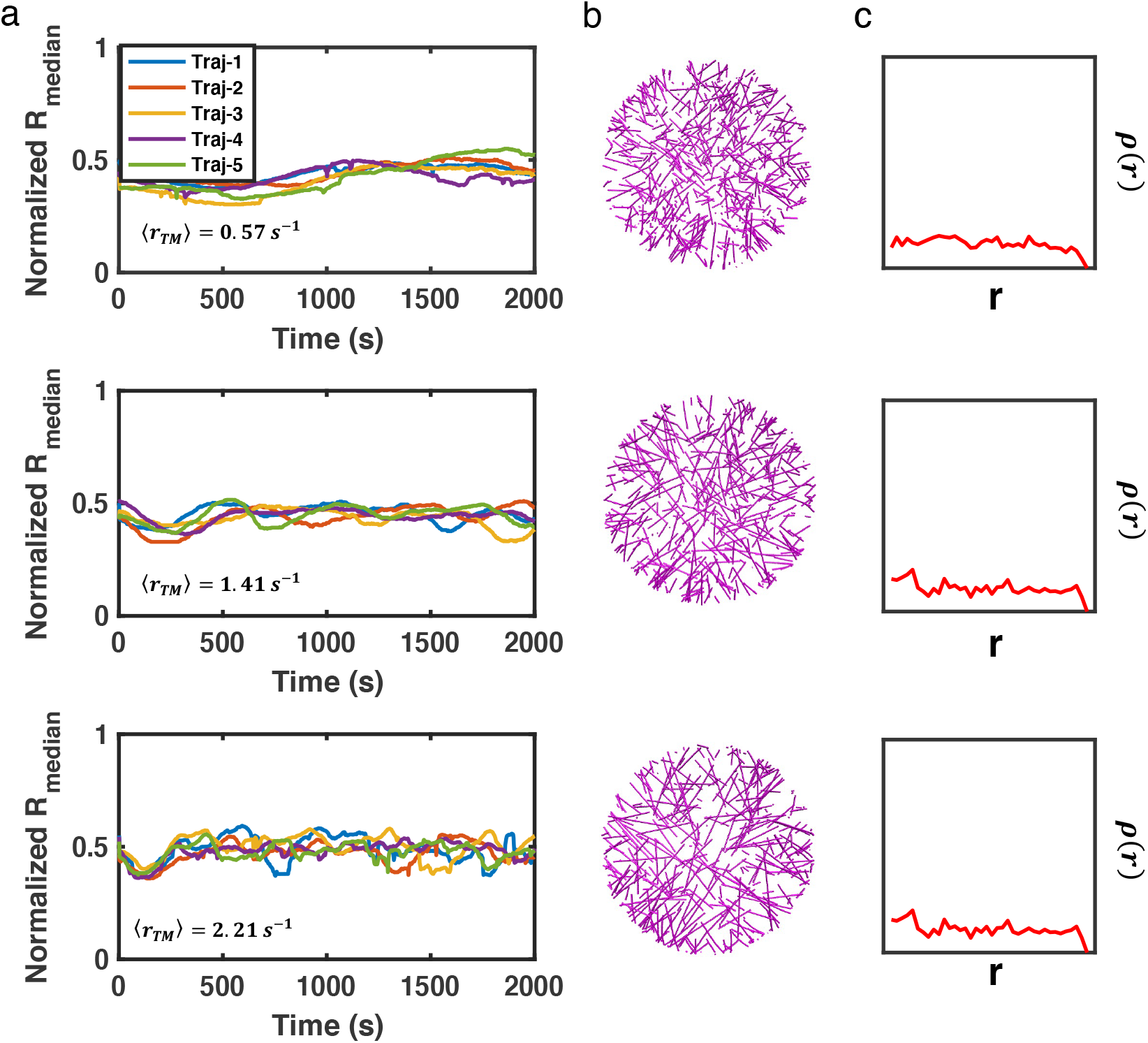
(a) 5 trajectories of normalized medians of filament radial density distribution, (b) representative snapshots, and (c) the corresponding filament radial density distributions (*ρ*_*r*_) at t = 2000 s are shown.

**Figure S4:**
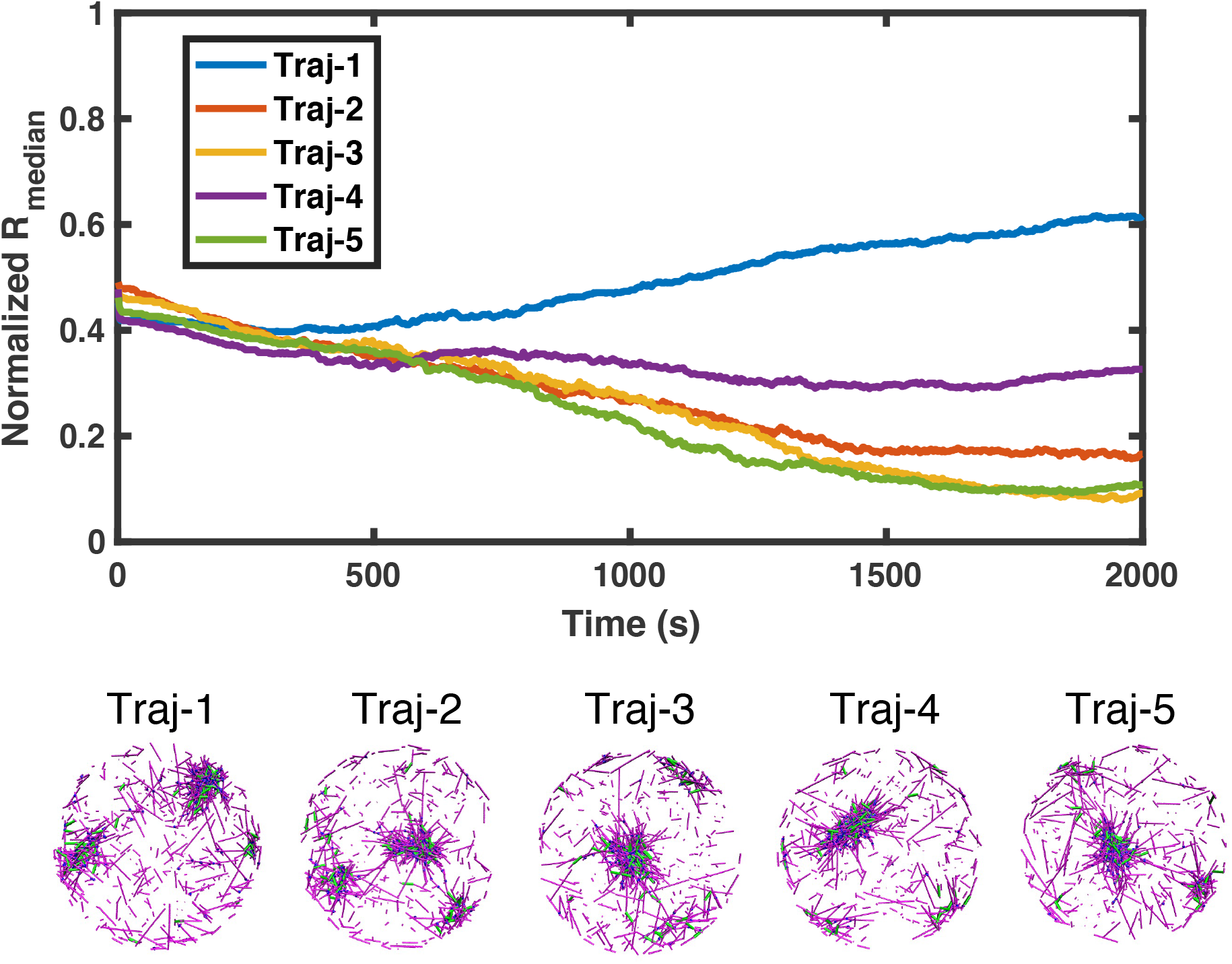
The upper plot shows 5 trajectories of medians of filament radial density distribution with *r*_*TM*_ = 0.56*s*^−1^. The lower part shows snapshots of each trajectory at the end of simulation.

**Figure S5:**
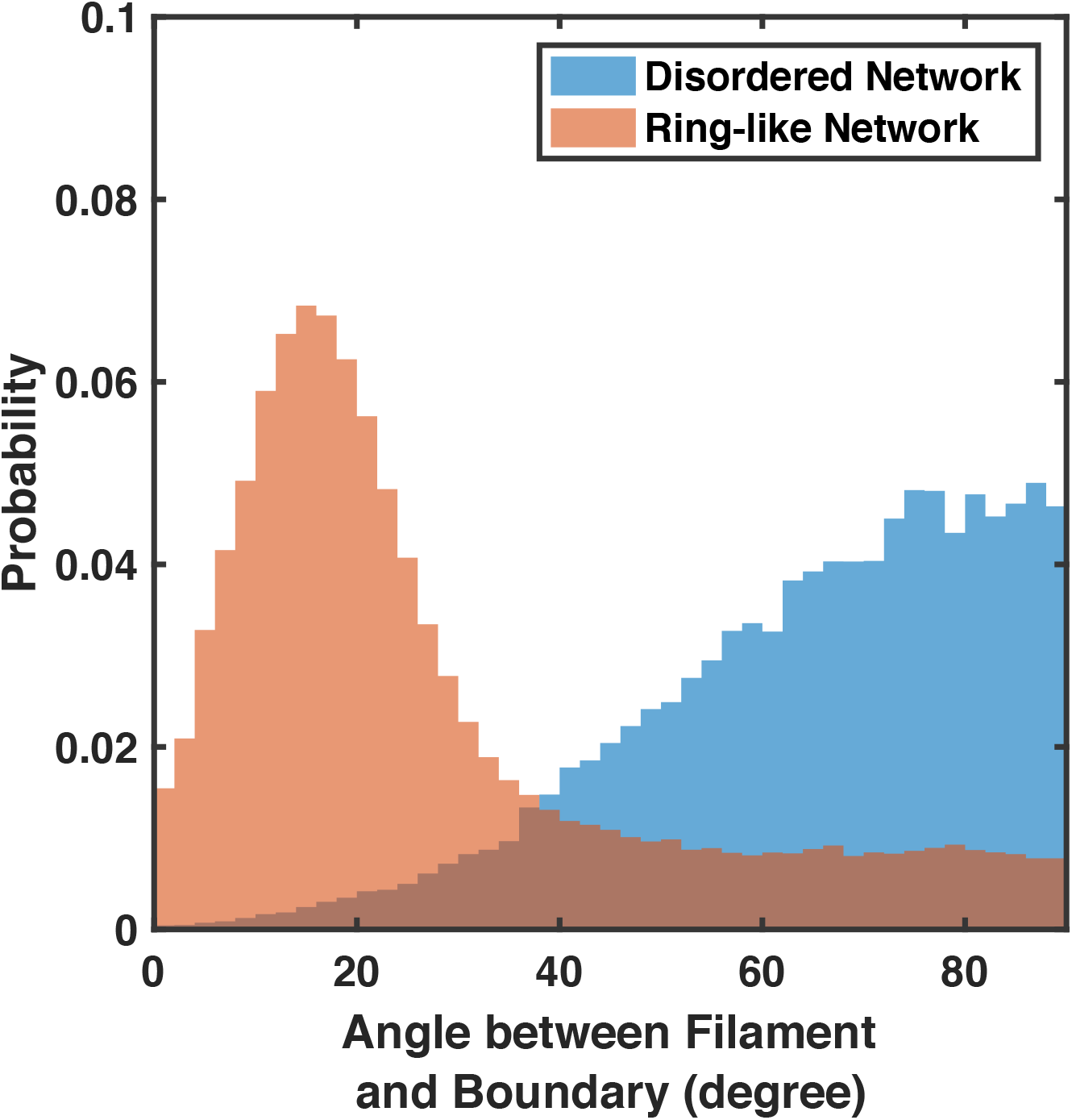
The distribution of filament orientations for disordered networks (⟨*r*_*TM*_⟩= 2.21*s*^−1^) and ring-like networks (⟨*r*_*TM*_⟩= 2.05*s*^−1^) near the network periphery (*r >* 1600*nm*) are shown. More filaments are oriented perpendicular to the boundary in disordered networks. The filament orientation is represented by the angle between the treadmilling direction (the non-bendable barbed end cylinder) and the tangent vector to the boundary. Only filaments longer than 200nm are counted. The angle ranges from 0° to 90°, with 0° indicating treadmilling parallel to the boundary and 90° indicating treadmilling perpendicular to the boundary. 5 runs per condition.

**Figure S6:**
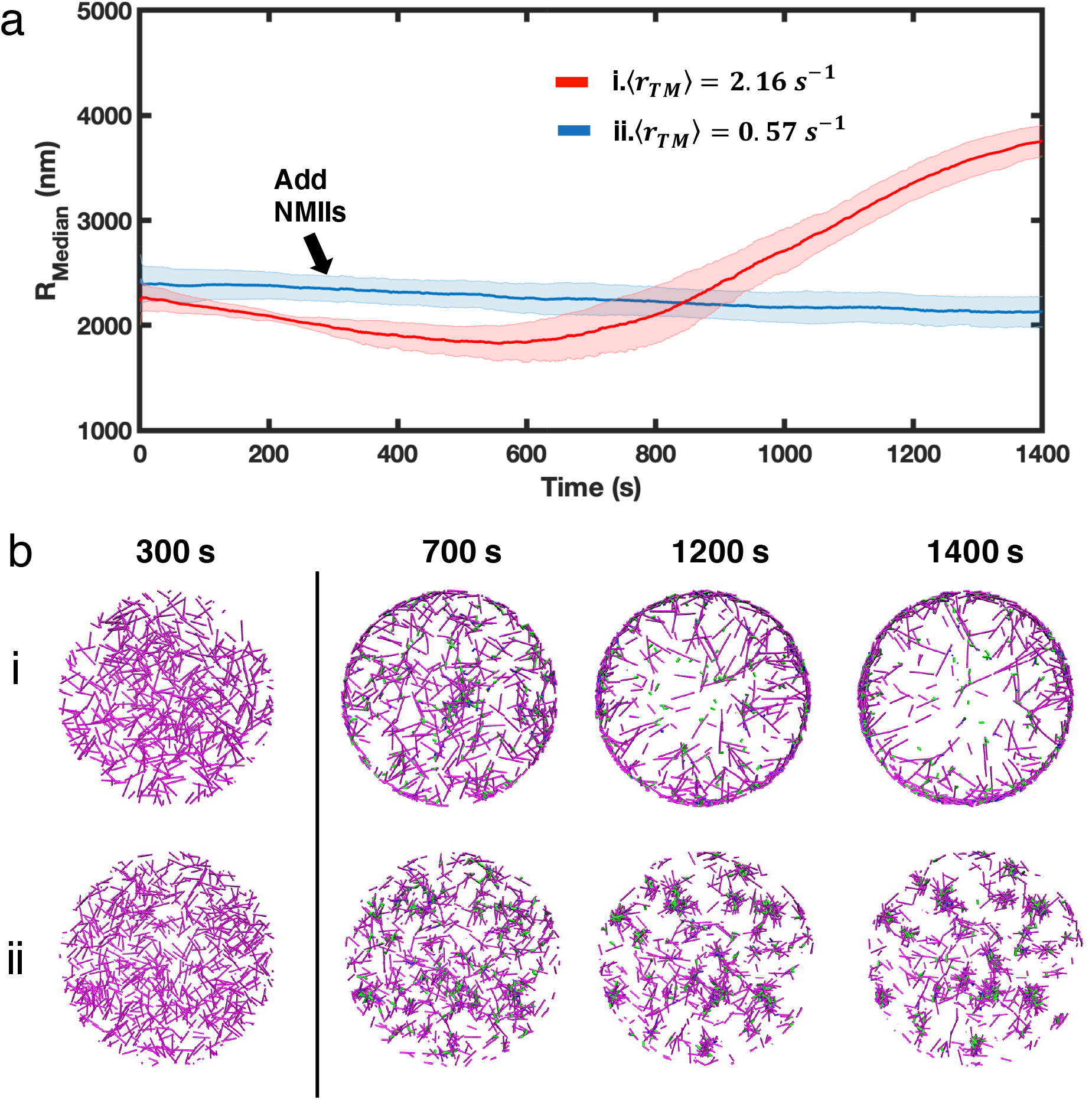
(a) Medians of filament radial density distribution (*R*_*median*_) at different treadmilling rates (⟨*r*_*TM*_⟩) are shown. Networks are thin oblate geometry, which has a diameter of 10 µm and a height of 200 nm. The network contains 20 µM G-actin and 30 nM filament nucleator. 0.03 µM of NMII and 2 µM alpha-actinin are added at t = 300 s.

**Figure S7:**
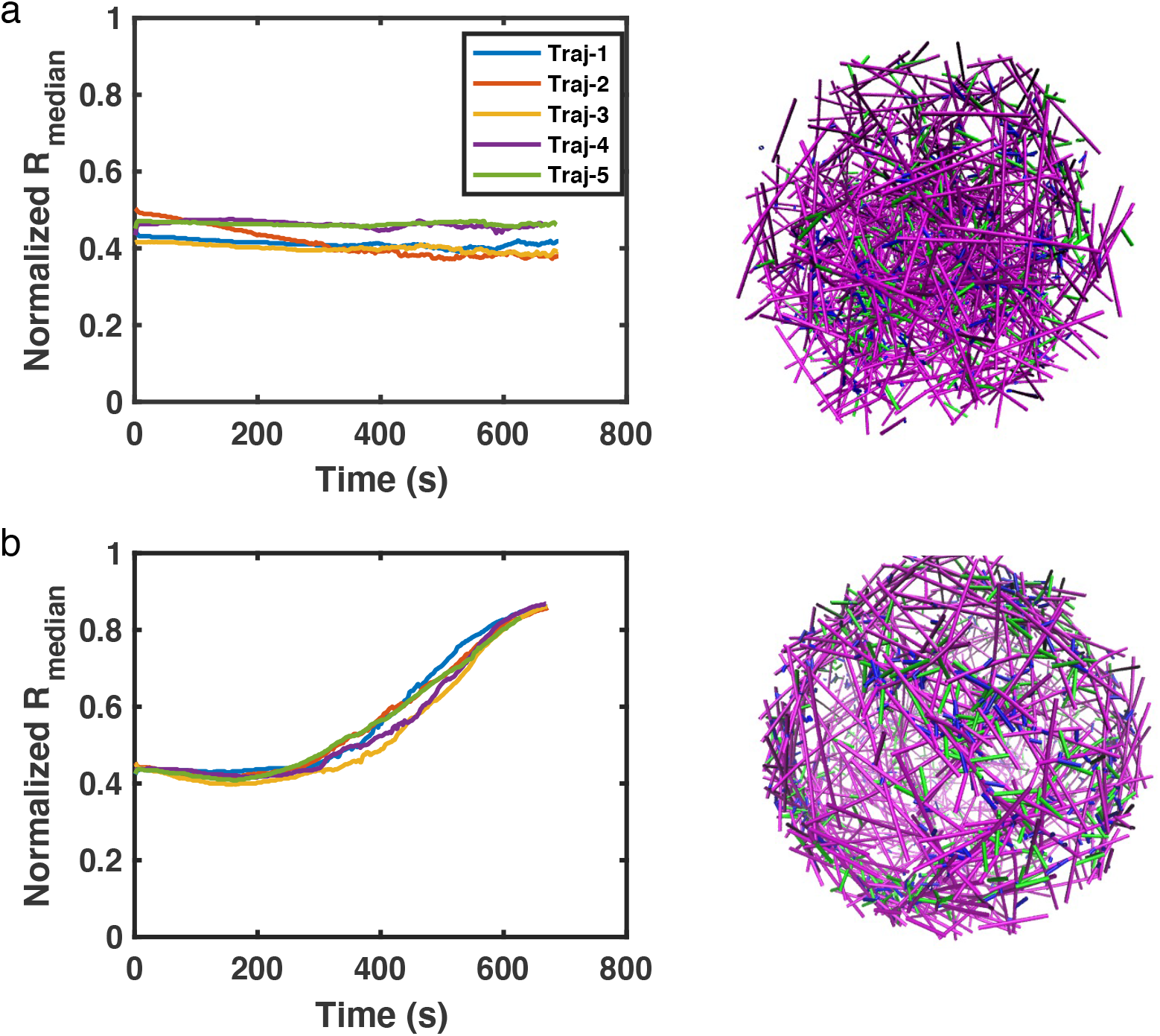
(a-b) Median of filament radial density distribution (*R*_*median*_) at different tread-milling rates (⟨*r*_*TM*_⟩) and the most representative snapshots in spherical networks are shown. The diameter of the spherical network is 4 µm. The network contains 20 µM G-actin and 20 nM filament nucleators. 0.03 µM of NMII and 2 µM alpha-actinin are added at t = 50 s.

**Figure S8:**
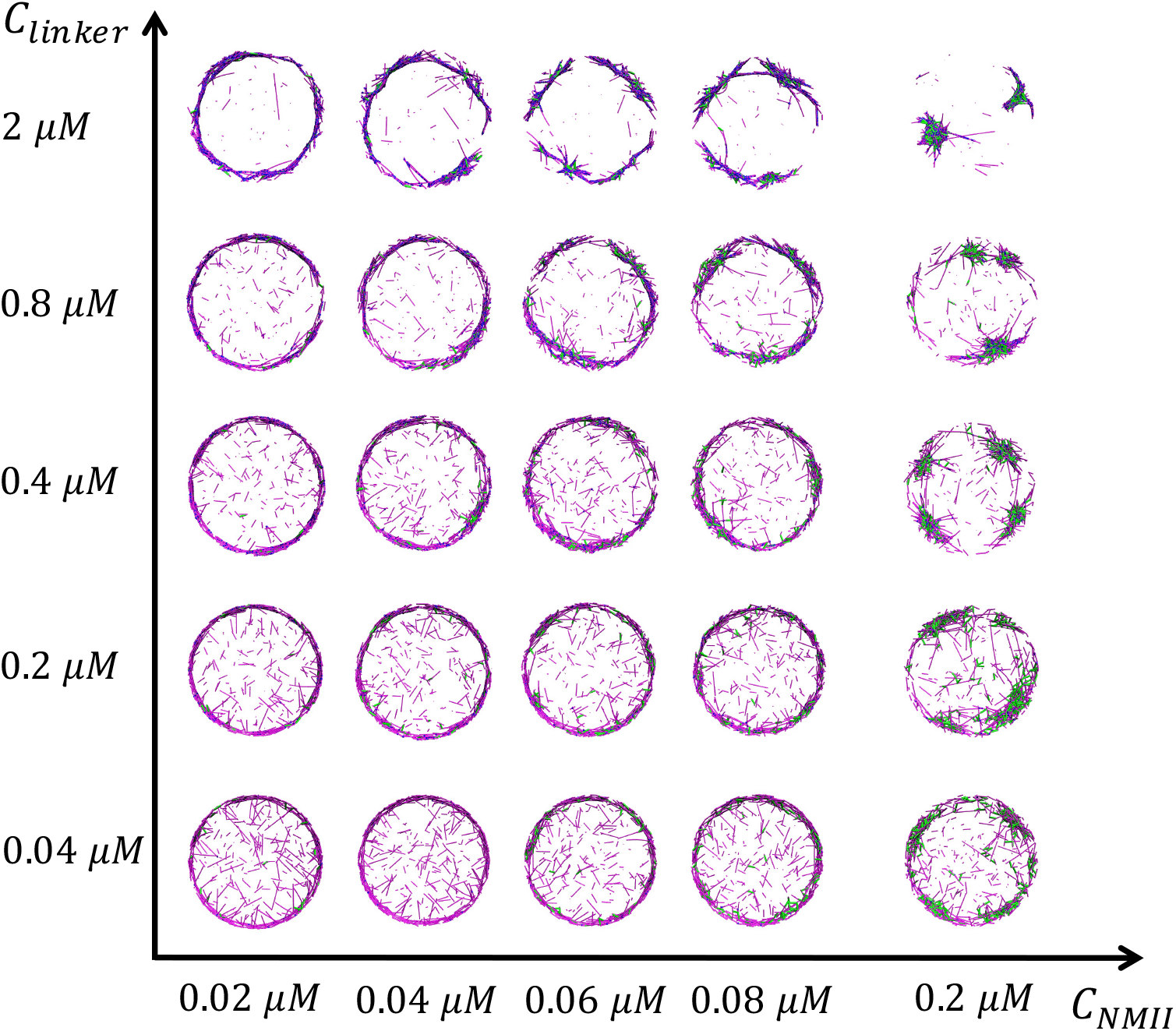
Representative snapshots of steady state actin network structures at different *C*_*alpha*−*actinin*_ and *C*_*NMII*_. ⟨*r*_*TM*_⟩ = 2.05*s*^−1^ for all conditions.

**Figure S9:**
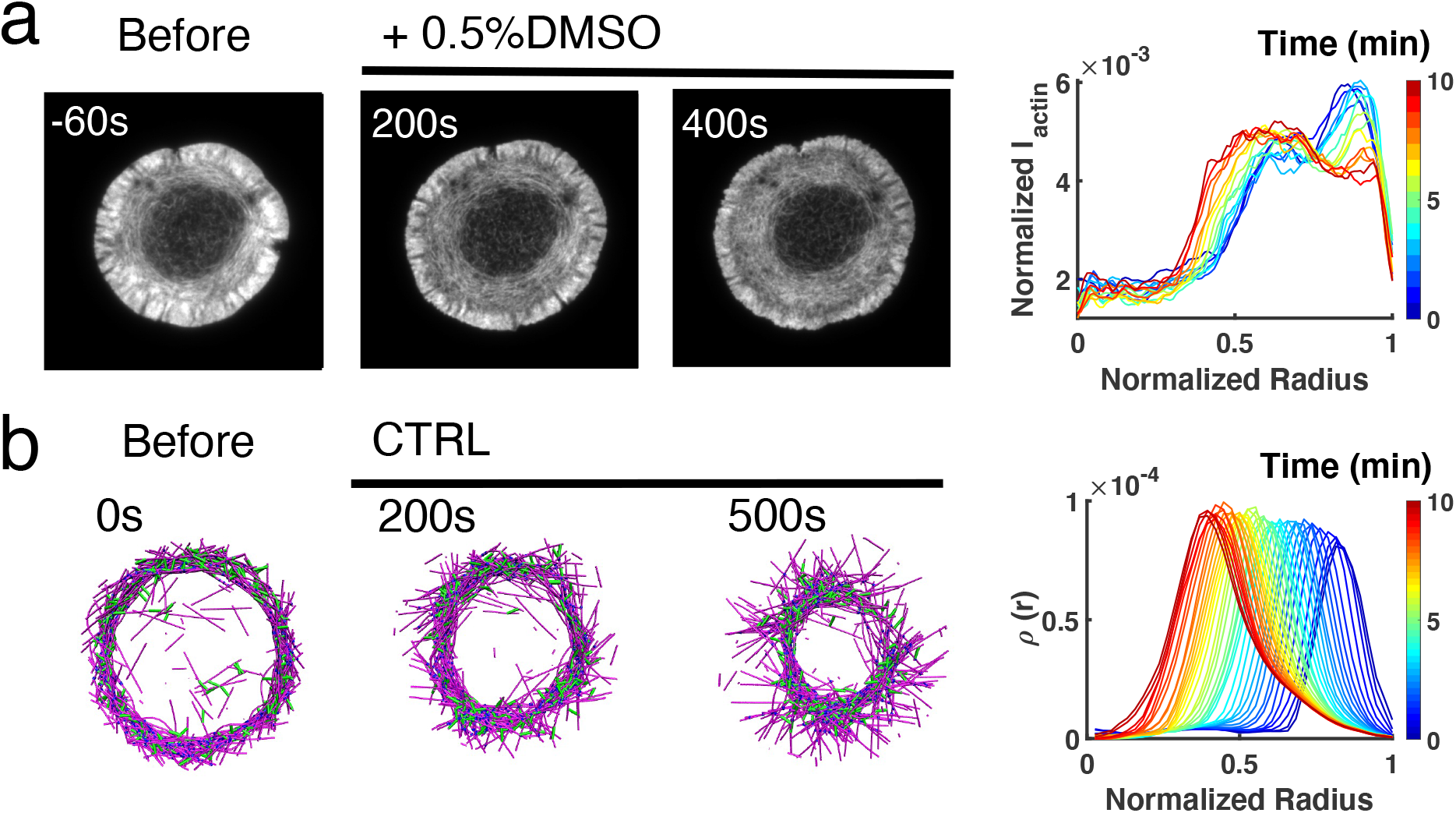
Enhancement or inhibition of NMII regulates actin structure in live T cells and *in silico*. (a-b) Time lapse montages of Jurkat T cells expressing F-tractin-EFGP spreading on anti-CD3 coated glass substrates (left) and the normalized radial filament density distribution *ρ*(*r*) at different time (right). After achieving maximal spreading, cells were treated with 0.5% DMSO. Scale bar is 10 µm. (b) Timelapse montages of simulations (left) and *ρ*(*r*) at different times (right) mimicking actin rings in vehicle control. Scale bar is 1 µm.

**Figure S10:**
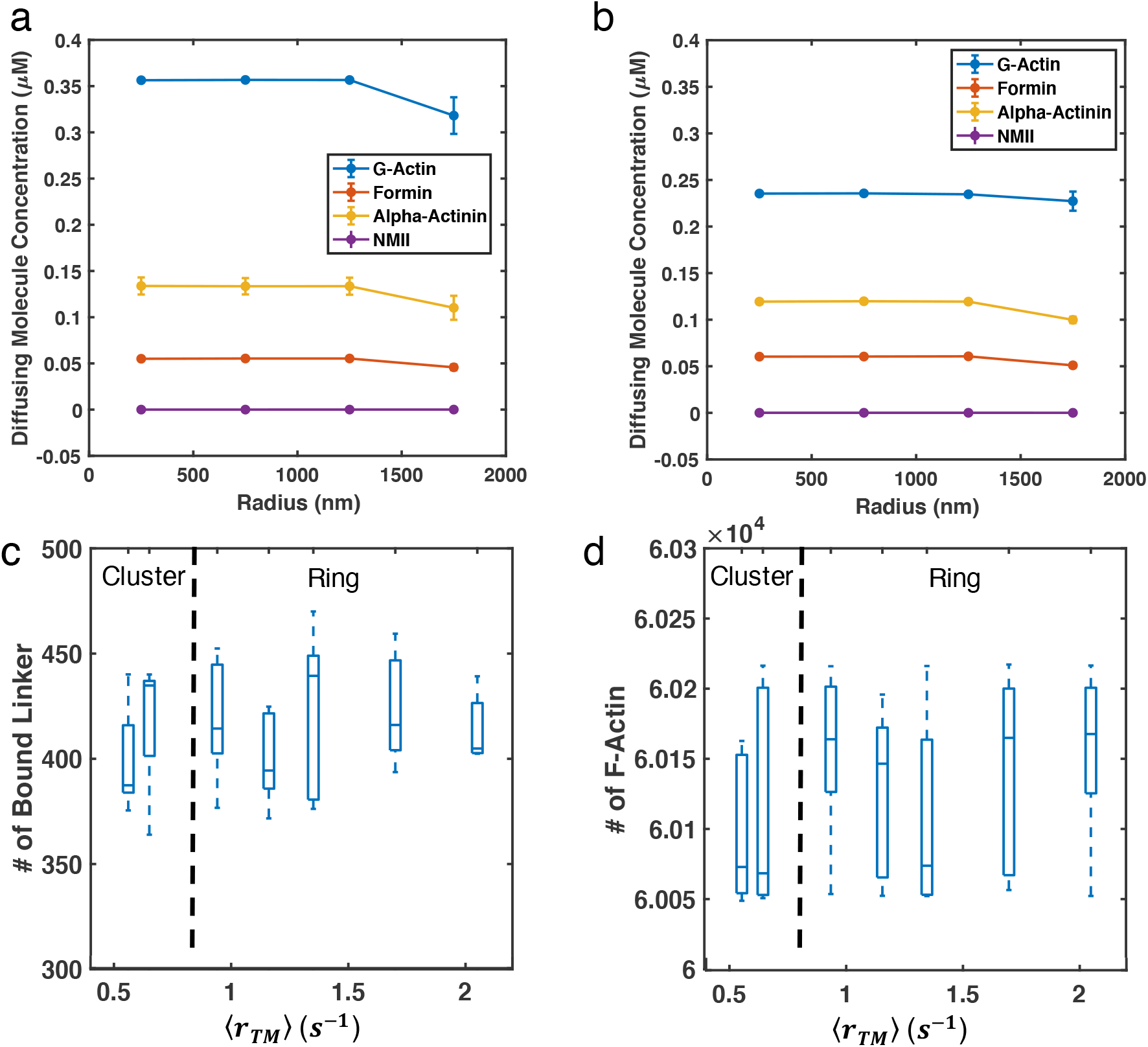
(a-b) Diffusing molecule concentrations along the radius of (a) cluster networks (⟨*r*_*TM*_⟩ = 0.56*s*^−1^) and (b) ring-like networks (⟨*r*_*TM*_⟩ = 2.05*s*^−1^, b) are shown. (c-d) Box plots of the number of (c) bound linker and (d) F-actin in the system are shown. Almost all motors are bound upon addition, thus we do not provide a plot for the number of bound motor.

## Supplementary Videos

Video 1. Timelapse movie of F-actin and NMII in Jurkat T cells activated by anti-CD3 coated stimulatory coverslips as shown in Figure 1a. F-actin is labeled by tdTomato-F-Tractin, and NMII is labeled by MLC-EGFP. Scale bar is 10 µm. Timestamp indicates time since seeding cells.

Video 2. The simulated actin network contains only actin filaments, as shown in Figure S3, and networks contain actin filaments, myosin, and crosslinkers with average tradmilling rate ⟨*r*_*TM*_⟩ = 0.56, 0.94, and 2.05*s*^−1^, respectively, as shown in Figure 2a-b. Actin filaments are magenta cylinders, and NMIIs are green cylinders.

Video 3. A zoomed in view of the fast-treadmilling network as shown in Video 3. During the actin ring formation, filaments (red cylinders) change their orientation from perpendicular to the boundary to in parallel to the boundary. Some filaments are highlighted with black to emphasize this deformation. NMIIs are blue cylinders.

Video 4. The evolution of a fast-treadmilling network in a spherical boundary as shown in Fig. 2e. The upper left part of the network is cropped to better illustrate the internal structure. Actin filaments are magenta cylinders, and NMIIs are green cylinders.

Video 5. Timelapse movie of F-tractin-EGFP labeled F-actin in Jurkat T cells activated by anti-CD3 coated stimulatory coverslips with 0.05% DMSO (vehicle control), and with 250nM, 500nM, and 1 µM LatA, respectively. Vehicle (DMSO) is added at 5 minutes. Scale bar is 10 µm. Timestamp indicates time since seeding cells.

Video 6. Simulation of disrupting actin assembly in ring-like actin networks that mimics LatA inhibition. The network is allowed to evolved for 800 seconds with ⟨*r*_*TM*_⟩ = 2.05*s*^−1^ as shown in video 2 before the disruption of actin filament polymerization and depolymerization. Actin filaments are magenta cylinders, and NMIIs are green cylinders.

Video 7. Timelapse movie of F-tractin-EGFP labeled F-actin in Jurkat T cells activated by anti-CD3 coated stimulatory coverslips with 50nM CalyA and 100 µM Y-27632, respectively. Scale bar is 10 µm. Timestamp indicates time since seeding cells.

Video 8. Simulation of hyper-activating and inhibiting NMIIs in ring-like actin networks that mimics CalyA and Y-27632 treatment, as shown in Figure 5. Networks were pre-assembled into actin rings during the initial 100s before hyper-activating and inhibiting NMIIs. Actin filaments are magenta cylinders, and NMIIs are green cylinders.

