## Supplemental Data 2 for "A tug of war between filament treadmilling and myosin induced contractility generates actin ring"

[1]Department of Chemical and Biomolecular Engineering
[2]Department of Physics
[3]Biological Sciences Graduate Program
[4]Biophysics Program
[5]Institute for Physical Science and Technology
[6]Department of Chemistry and Biochemistry
University of Maryland, College Park, MD, USA

[†]These authors contribute equally.

#### 1 Computational Model Details

The simulations presented in this work were carried out via MEDYAN (Mechanochemical Dynamics of Active Networks). MEDYAN is an open-access mechanochemical simulation platform (available at www.medyan.org) for active matter (*1*). We employed MEDYAN to model stochastic reaction-diffusion, mechanical interaction, and mechanochemical dynamics of active cytoskeletal networks.

In MEDYAN, the simulation space is divided into a solution phase and a polymer phase. All diffusing molecules (DM) are dissolved in the solution phase, including G-actin monomers, unbound formin, unbound non-muscle myosin II (NMII), and unbound crosslinker molecules. To reduce the computational cost without losing too much spatial information, the solution phase is discretized into many linear compartments. The dimension of the compartment is carefully chosen based on the Kuramoto length of actin, which is the mean-free path that molecules are expected to diffuse before undergoing their next reaction (*2*). In the spherical network, compartments are 500 nm $\times$ 500 nm $\times$ 500 nm cubes. In thin oblate networks, the compartment dimension in the Z-axis is reduced to $400nm$ (the mechanical boundary is $200nm$ as restricted by the boundary repulsion potential, see Section 1.2 for details). Diffusing molecules are assumed to be well-mixed within each compartment without specific spatial locations, and the transport of molecules between compartments is modeled as a diffusion reaction within the solution phase.

On the other hand, polymeric filaments, bound formin, bound NMIIs, and bound crosslinkers comprise the polymer phase that lays over the solution phase. This phase accounts for the mechanical modeling of boundary repulsion, steric interactions, bending and stretching of filaments, as well as the stretching of linkers and motors. When polymerization, nucleation, NMII or crosslinker binding reactions occur, diffusing molecules transfer from the solution phase into the polymer phase. Depolymerization, filament destruction, NMII or crosslinker unbinding reactions will release molecules from the polymer phase to solution phase.

In the following sections, we will further discuss details of our mechanical models, chemical reaction-diffusion models, mechanochemical coupling, and the simulation protocol.

#### 1.1 Mechanical models

Unlike the traditional bead-spring model, the semi-flexible filaments are represented as connected cylinders. The equilibrium length (under zero force) of each cylinder elements varies from 2.7nm (1 actin monomer) to a maximum of 108nm (40 actin monomers). Addition of each actin monomer would increase the length of the first or last cylinders by 2.7nm, and vice versa. Polymerization will create a new cylinder if the cylinder has reached its maximum length. Filaments have a very large aspect ratio, i.e., the persistence length of a filament ($\sim 20\mu m$) is much larger than its diameter ($\sim 10nm$). Thus, it is reasonable to ignore the radial stretching/compression and only allow the axial stretching/compression of a cylinder, which is written as

$$U^{str}_{filament} = \frac{1}{2} K^{str}_{filament} (l_f - l_{f,0})^2.$$

$l_f$ is the actual length of cylinder under force, and $l_{f,0}$ is the equilibrium length based on the number of actin monomers on this cylinder (each monomer is 2.7nm). Radial filament deformation is modeled as bending between two connected cylinders:

$$U^{bending}_{filament} = K^{bending}_{filament} (1 - cos(\theta - \theta_0)),$$

where $\theta$ is the angle between the two consecutive cylinders under force, while $\theta_0$ is the equilibrium angle that is set to be 0.

A novel volume exclusion exclusion potential is implemented to prevent cylinders overlapping, which is written as

$$U^{Vol} = \iint_{l_i, l_j} \delta U \mid \vec{r_i} - \vec{r_j} \mid dl_i dl_j,$$

where $\delta U \mid \vec{r_i} - \vec{r_j} \mid = 1 / \mid \vec{r_i} - \vec{r_j} \mid^4$ is the pair potential between two points located on the two interacting cylinders. $\vec{r_i}$ amd $\vec{r_j}$ are the distances between any two points along the cylinder $i$ and $j$, respectively. This potential can provide a steep enough volume exclusion effect while remain analytically solvable.

Bound NMIIs and linkers are modeled as harmonic springs, and the stretching energy is written as

$$U^{str}_{NMII/linker} = \frac{1}{2} K^{str}_{NMII/linker} (l_{NMII/linker} - l_{NMII/linker,0})^2.$$

Table SI-1: Mechanical parameters.

| Names | Parameters | References |
|---|---|---|
| Cylinder stretching | $K^{str}_{filament} = 100pN/nm$ | 1 |
| Cylinder bending | $K^{bending}_{filament} = 672pN \cdot nm$ | 3 |
| Filament volume exclusion | $K_{vol} = 10^5 pN/nm^4$ | 1 |
| Linker stretching | $K^{str}_{linker} = 8pN/nm$ | 4 |
| NMII stretching | $K^{str}_{NMII} = 2.5pN/nm$ per head | 5 |
| Boundary repulsion | $\epsilon_{boundary} = 100pN \cdot nm$ | This work |

$l_{NMII/linker,0}$ is the equilibrium length of a linker, which are initialized when a linker /NMII binding reaction occurs as the distance between the paired binding site. $l_{NMII,0}$ is reset every time a motor walking reaction occurs.

In order to confine all the filaments within the simulation boundary, an exponential boundary repulsion potential is implemented. In the thin oblate system, the actual height of the network is set to be 400 nm, and the diameter to 4000 nm. However, filaments would occasionally move out of the mechanical boundary due to rapid treadmilling, leading to simulation failures. To prevent this, we shift the boundary barrier slightly inside the network by $a_0$, and the exponential boundary repulsion is written as

$$U^{boundary} = \epsilon_{boundary} e^{-(d-a_0)/\lambda},$$

where $\epsilon_{boundary} = 100pN \cdot nm$ is the repulsive energy constant, $d$ is the distance between boundary and filament element, and $\lambda = 2.7nm$ is the screening length. The boundary shifting factor $a_0$ is chosen to be 100 nm based on experience. The existence of $a_0$ restricts the effective network boundary to height = 200 nm and diameter = 3800 nm.

The mechanical model parameters can be found in Table SI-1.

#### 1.2 Chemical models

The chemical engine of MEDYAN is powered by Next Reaction Method (NRM) (*6*), which is a variant of the Gillespie algorithm (*7*). Overall, the NRM stochastically solves the chemical Master Equation by generating a trajectory of chemical events. In this work, we simulated the following chemical reactions: diffusion, filament polymerization, filament depolymerization, filament nucleation, destruction of filaments, binding of myosin motors and linkers, and motor walking.

The diffusion of molecules is modeled as a single molecule transfer process between neighboring compartments, which follows our stochastic chemical reaction protocol as

$$DM_{i_0,j_0,k_0} \longrightarrow DM_{i_1,j_1,k_1},$$

where a diffusing molecule (DM) originally located in compartment $_{i_0,j_0,k_0}$ is transferred to a neighboring compartment $_{i_1,j_1,k_1}$. The copy number of this diffusing molecule species is decreased by 1 in compartment $_{i_0,j_0,k_0}$ and is increased by 1 in compartment $_{i_1,j_1,k_1}$.

Actin filament (F-actin) polymerization and depolymerization occur at both barbed end (BE) and pointed end (PE) of a filament. These reactions are written as

$$G - actin \longrightarrow F - actin,$$

$$F - actin \longrightarrow G - actin$$

It should be noted that G-actin is dissolved in the solution phase, while F-actin is in the polymeric phase.

The nucleation reaction is presented as a two-step reaction based on the mechanism of formin nucleation (*12, 13*):

$$Step\,1 : Formin + G - actin \longrightarrow intermediate,$$

Table SI-2: Parameters for diffusion and reactions.

| Names | Parameters | References |
|---|---|---|
| Diffusion | $D_{actin,arp2/3,CP} = 20\mu M^2/s$ | 2 |
| Actin | $k_{on}^{BE} = 11.6 - 34.8(\mu M \cdot s)^{-1}$ | 8 and this work |
| | $k_{on}^{PE} = 1.3(\mu M \cdot s)^{-1}$ | |
| | $k_{off}^{BE} = 1.4s^{-1}$ | |
| | $k_{off}^{PE} = 0.8 - 2.4s^{-1}$ | |
| Destruction | $k_{destruction} = 1.0 - 1.9s^{-1}$ | This work |
| Nucleation | $k_{nu} = 0.005s^{-1}$ | [Turnover] |
| Formin dissociation | $k_{off}^{formin} = 0.01s^{-1}$ | 9 |
| Alpha-actinin | $k_{on}^{\alpha} = 0.7(\mu M \cdot s)^{-1}$ | 10 |
| | $k_{on}^{\alpha} = 0.3s^{-1}$ | |
| NMII head binding | $k_{on}^{M} = 0.2s^{-1}$ | 11 |
| | $k_{on}^{M} = 1.7s^{-1}$ | 1 |

$$Step\ 2 : G - actin + intermediate \longrightarrow FBE - actin + F - actin + PE - actin.$$

The intermediate is an arbitrary molecule that consists of a formin and a G-actin molecule. We assume step 1 is the rate-limiting step and step 2 is a fast step, thus this intermediate would rapidly react with a G-actin molecule and become a short filament consisting of one F-actin molecule at the pointed end (PE-actin), a regular F-actin molecule, and another F-actin molecule at the formin bound barbed end (FBE-actin). For simplicity, polymerization and depolymerization at FBE are the same as regular barbed end reactions. Formin can dissociate from a filament, which releases a formin molecule into the solution phase and creates a regular F-actin barbed end (BE-actin) on that filament:

$$F - actin + FBE - actin \longrightarrow BE - actin + Formin.$$

Since new filaments are constantly created by nucleation,, the filament destruction process is required to establish a steady state which maintains a constant total number of filaments. The destruction reaction occurs exclusively when a filament has only two F-actin molecules (a BE-actin and a PE-actin), which destroys this filament and releases two diffusing G-actin molecules as

$$BE - actin + PE - actin \longrightarrow 2G - actin.$$

The binding reactions of myosin motors and linkers are carried out with a slightly different protocol. Firstly, the system will search for all possible binding site pairs on actin filaments and stochastically choose one for binding reaction. The two binding sites of a pair must be located at different filaments. The distance between the two binding sites ranges from 175-225 nm for NMII mini filament (*14*), and 30-40 nm for alpha-actinin crosslinker (*15*). After the binding site pair is determined, the binding reaction convert a diffusing motor or linker to a bound motor or linker with two ends attaching to the two binding sites, creating a mechanical linkage. This linkage vanishes when an unbinding reaction occurs, releasing the motor or linker to the diffusing pool. It should be noted that NMII mini filament is an ensemble of 15-30 myosin heads (*16*), and we model the entire ensemble as a while. To take the variation of the number of myosin heads into account, the number of myosin heads of each NMII mini filament is chosen stochastically for each reaction, and the reaction rate for each NMII binding event is then scaled by the number of myosin heads. .

In an active cytoskeleton, myosin motors consume energy from ATP hydrolysis and actively walk along filaments, which is one of the most important sources of contractile force generation. In MEDYAN, a motor stepping reaction is implemented to mimic this effect. For a bound NMII, the stepping reaction is written as

$$NMII_i \longrightarrow NMII_{i+1},$$

where $i$ and $i + 1$ are the NMII locations on the filament before and after walking. NMII is a barbed end walking motor, thus $i + 1$ represents the next binding site towards the barbed end.

Parameters for diffusion and chemical reactions can be found in Table SI-2.

#### 1.3 Mechanochemical models

Many cytoskeletal reactions, including actin polymerization, myosin motor binding and stepping, and linker binding, are mechanosensitive. To capture this feature, MEDYAN implements mechanochemical models that explicitly allow force-dependent chemical reaction rates.

Table SI-3: Mechanochemical dynamic rate parameters.

| Names | Parameters | References |
|---|---|---|
| Characteristic polymerization force | $F_{poly,0} = 1.5pN$ | 17 |
| Characteristic linker unbinding force | $F_{linker,unbind} = 17.2pN$ | 18 |
| NMII duty ratio | $\rho = 0.1$ | 11 |
| NMII stall force | $F_{stall} = 12.62pN$ per head | 19 |
| Tunable parameters | $\beta = 0.2$ | 1 |
| | $\gamma = 0.05pN^{-1}$ | |
| | $\xi = 0.1$ | |

The effect of boundary force on filament polymerization is described by the Brownian Ratchet model (*20*), which models the force sensitive polymerization rate $k_{poly}$ as:

$$k_{poly} = k^0_{poly} \cdot \exp(-F_{ext}/F_{poly,0}),$$

where $k^0_{poly}$ is the bare polymerization rate under zero external force, $F_{ext}$ is the boundary repulsive force exerted on the filament ends, and $F_{poly,0}$ is the characteristic polymerization force based on the thermal energy and the size of actin monomers.

We used a simple exponential equation to model the slip bond property of alpha-actinin crosslinker:

$$k_{linker,unbind} = k^0_{linker,unbind} \cdot \exp(F_{linker,stretching}/F_{linker,unbind}),$$

where $k^0_{linker,unbind}$ is the unbinding rate constant under zero external force, and $F_{linker,unbind}$ is the characteristic unbinding force of alpha-actinin. $F_{linker,stretching}$ is the stretching force on the linker, while a compressive force on the linker does not trigger the slip bond.

In this work, we model NMII binding as a catch bond, as adapted from the Parallel Cluster Model (*19*), such that the force loaded on NMII can reduce its unbinding rate constant:

$$k_{NMII,unbind} = \frac{\beta \cdot k^0_{NMII,unbind}}{N_{heads}} \cdot exp(\frac{-F_{ext}}{N_{heads} \cdot F_{NMII,unbind}}),$$

where $\beta$ is a tunable parameter, $k^0_{NMII,unbind}$ is the unbinding rate constant under zero force, $F_{ext}$ is the total stretching force applied on the NMII, and $N_{heads}$ is the number of NMII heads.

The NMII walking rate is also mechanochemically sensitive and can be modeled with a Hill type force-velocity relation:

$$k_{walk} = k^0_{walk} \cdot \frac{F_{stall} - F_{ext}/N_{heads}}{F_{stall} + F_{NMII,pulling}/(N_{heads} \cdot \xi)},$$

where $F_{stall}$ is the stall force of a single NMII head, $F_{NMII,pulling}$ is the pulling force on NMII in the opposite direction of walking movement, and $\xi$ is a tunable parameter.

The mechanochemical model parameters can be found in Table SI-3.

#### 1.4 Simulation protocol

The relaxation time for local deformations of actin networks (*21*) is much shorter than the timescale of typical chemical events such as motor stepping (*11*) or filament polymerization (*8*), thereby creating a significant separation of timescales. Hence, the mechanical equilibrium process can be viewed as a pseudo-adiabatic process that can be separated from chemical reactions. Based on this hypothesis, the simulation can be carried out in the following steps:

1. Chemical reactions occur that evolve the time of the system stochastically.

2. Pausing chemical reactions when the time step reaches a preset value, which is 10 ms in this work. The system then mechanically minimizes the total energy.

3. Reaction rates are updated based on the tension acting on NMIIs/linkers and load force acting on actin filament barbed ends after mechanical minimization.

4. Step 1 is repeated based on the updated reaction rates.

This protocol is iterated until we reach 2000 seconds of simulation time, or until we reach the wall time limit on the Deepthought2 High-Performance Computing cluster at University of Maryland, College Park, whichever comes first.

### 2 Defining treadmilling rate and treadmilling inhibition simulation setups

Although treadmilling in cells is a complex system that involves hundreds of reactions (*22, 23*), it is simplified to four reactions in this work by considering polymerization and depolymerization at both barbed ends and pointed ends. When a steady state is established, the net barbed end growth rate will equal the net pointed ends reduction rate (averaged over the system), maintaining a constant average filament length. Therefore, we can define a kinetic steady state for treadmilling by monitoring the average filament length of the network as shown in Fig. SI-1. We found that such a kinetic steady state could be established after 1000s in all conditions, and at this state, the average barbed end elongation rate is almost the same as the average pointed end shrinkage rate. Hence, we quantify the average treadmilling rate $\langle r_{TM} \rangle$ as the average barbed end elongation rate after 1000s.

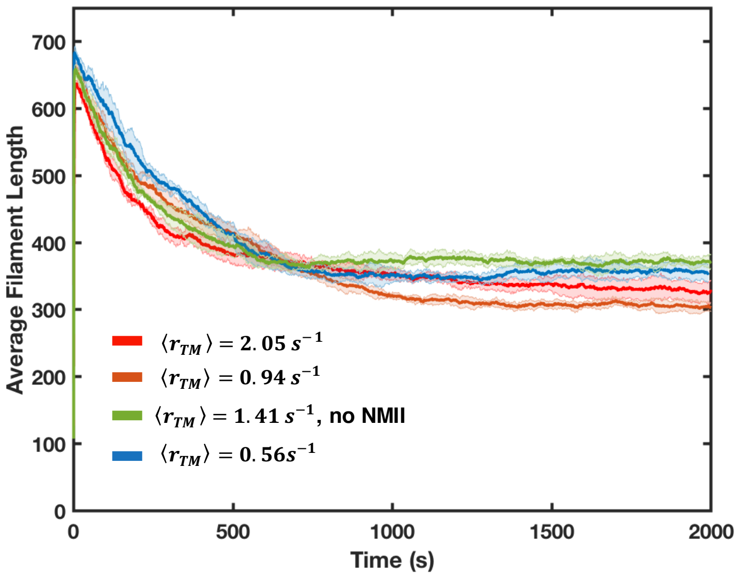


Figure SI-1: Average filament length as a function of time are shown. Shaded color represents the standard deviation of mean (5 run per condition).

While the treadmilling rate is an elegant and robust way of quantifying the speed of actin network assembly, it is extremely hard to measure *in vivo*. An alternative way to quantify the speed of actin network remodeling is to measure the turnover timescale, which has been widely studied via an experimental technique called Fluorescent Recovery After Photobleaching (FRAP). To compare with experiments, in our simulation we used a method mimicking the FRAP to calculate the turnover halftime ($t_{1/2}$, the time required for a network to reach 50% turnover) as developed in our previous work (*13*) , and we obtain $t_{1/2} \sim 168s$ for the slowest treadmilling condition, and $t_{1/2} \sim 48s$ for the most rapid treadmilling case. It should be noted that our longest $t_{1/2}$ is similar to the turnover timescale of some reconstituted networks (*24*), and our shortest $t_{1/2}$ is comparable to that of *in vivo* actin cortices (*25*). The details of turnover halftime measurement in MEDYAN and how it is related to treadmilling has been discussed in depth in a prior computational study (*13*).

We utilized kinetic parameters measured *in vitro* (*8*) as the baseline to assemble the slow treadmilling networks. To explore suitable parameters for rapidly treadmilling networks, we looked into the effects of formin and ADF/cofilin. An earlier work (*26*) has shown that the presence of formin can boost the polymerization rate at the barbed end several-fold over the baseline. For simplicity, we imitated this effect by increasing the barbed end polymerization rate constant ($k_{on}^{BE}$). ADF/cofilin can also promote treadmilling by severing filaments. Importantly, the fragment that contains the pre-existing pointed end is very unstable and would undergoes rapid disassembly (*24*). This observation allows us to mimic the effect of ADF/cofilin by simply increasing the depolymerization at the pointed end ($k_{off}^{PE}$). For example, we increase the $k_{on}^{BE}$ and $k_{off}^{PE}$ to three-fold in the actin ring network as shown in Fig. 1a-c ($\langle r_{TM} \rangle = 2.05 s^{-1}$).

### 3 Calculation of local actin concentration for clusters and rings

In this work, we used a density-based clustering method to define regions that contain actin clusters and rings, and calculated the local F-actin concentration within these regions. We first a generated pixelated map by dividing the network into $100nm \times 100nm$ bins and calculated the F-actin concentration within each bins (Fig. SI-2a). We then grouped connecting bins with concentration higher than a threshold (160 µM) into clusters (Fig. SI-2b). Clusters with size less than 4 bins were ignored. The local actin concentration within clusters was calculated as the average F-actin concentration of these clusters. The local actin concentration within actin rings is calculated using the same method (Fig. SI-2c-d).

### 4 Simulation setups of Latrunculin A, Calyculin A, and Y-27632 modeling

Earlier works have shown that LatA affects filament treadmilling in two ways: 1) it sequesters G-actin and 2) it accelerates the phosphate release from ADP-Pi-actin thereby reducing fila-

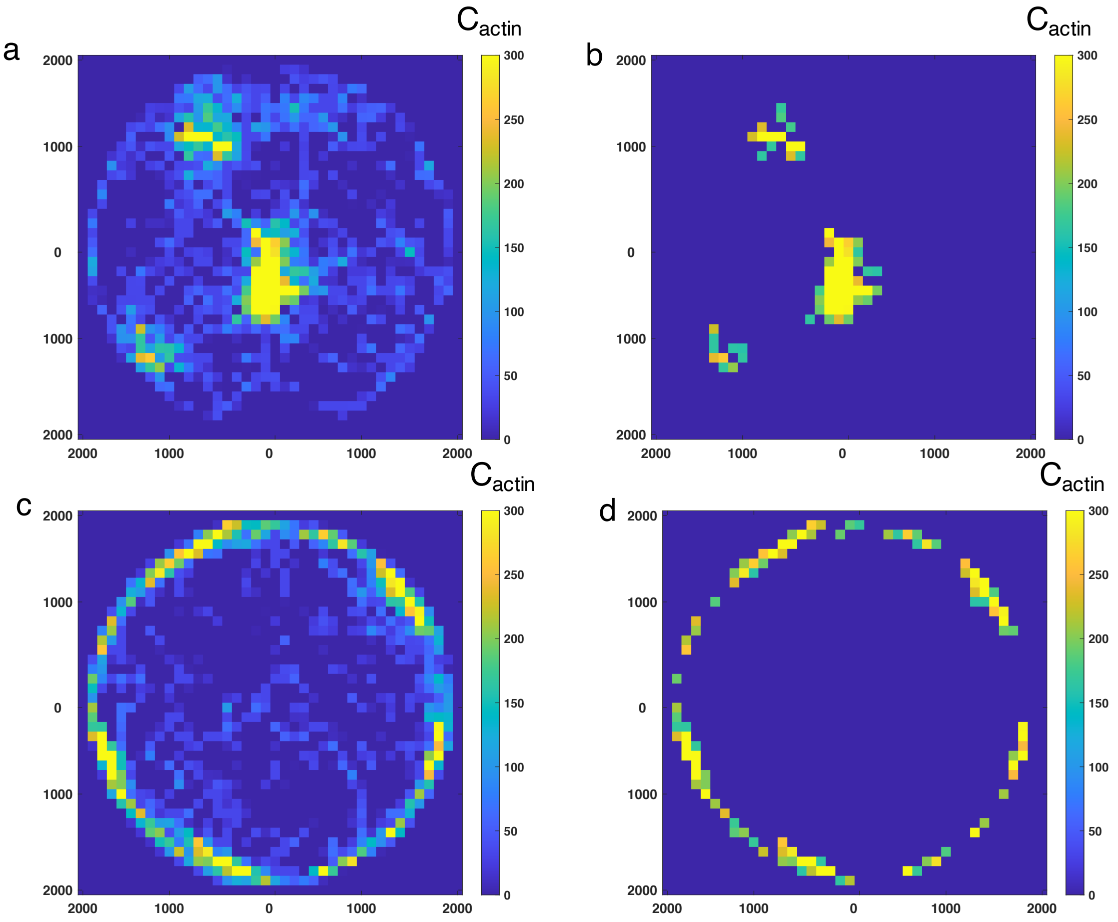


Figure SI-2: Heatmap showing a pixellated representation of the local F-actin concentration in 100nm x 100nm bins for (a) a cluster-like network as shown in Fig.1a-c ($\langle r_{TM} \rangle = 0.56s^{-1}$), and (b) a ring-like network as shown in Fig.1a-c ($\langle r_{TM} \rangle = 2.05s^{-1}$). (b and d) are the same heatmaps as (a) and (c), respectively, but only contains bins that exceed a threshold concentration of 160 µM.

ment polymerization while increasing depolymerization at both ends (*27–29*). To simulate such effects in the actin ring perturbation simulations, we explore a parameter space that mimicked the effect of LatA treatment: we disrupted $r_{TM}$ by reducing the filament polymerization rate and increasing the depolymerization rates. In the weak inhibition case, we decreased $k_{on}^{BE}$ to $11.6(\mu M \cdot s)^{-1}$, increased $k_{off}^{BE}$ to $2.1s^{-1}$, and maintained $k_{off}^{PE}$ at $2.4s^{-1}$. In the strong inhibition case, $k_{on}^{BE}$ was decreased to $3.48(\mu M \cdot s)^{-1}$, $k_{off}^{BE}$ was increased to $11.2s^{-1}$, and $k_{off}^{PE}$ was increased to $4.8s^{-1}$. In all simulations, pointed end polymerization rate was set to be constant at $1.3(\mu M \cdot s)^{-1}$. Treadmilling rate is consequentially reduced as a result of such disruption.

Calyculin A is an enhancer of NMII activity by inhibiting myosin light chain ATPase, while Y-27632 inhibits Rho kinase, a upstream regulator of NMII. Thus, we model their effects by increasing or decreasing the NMII levels after actin ring formation to match the T cell experiment. In the CalyA experiment, actomyosin ring collapses while maintaining the ring-like geometry. We realize that such "whole ring contraction" is difficult to achieve at the low actin concentration ($C_{actin} = 40\mu M$) that we used other conditions. At low actin concentration, enhancing NMII activity often simultaneously cause centripetal collapse as well as the local collapse that disassemble the ring-like structure, due to lack of filament-filament connectivity. To overcome this issue, we double the actin concentration to $C_{actin} = 80\mu M$ and adjust $C_{NMII}$ to 0.18 µM in the model. Such a high concentration of actin and motor protein significantly reduces the computational efficiency, therefore we initialize the ring-like actin structure instead of starting from a disordered network. At control condition as shown in Figure S9, network will slightly contract but can maintain the ring-like structure.

### 5 Calculating the slope of center to plateau F-actin distribution

After forming a cell mask (Fig. SI-3,left - yellow outline) using a minimum threshold intensity and identifying the centroid (red dot) of the masked cell, 50 lines joining the centroid to the mask edge are randomly drawn and the intensity profile averaged over all these lines is plotted. This plot gives a single intensity line profile from cell centroid to cell edge for a cell at a time point. Similarly, the line profiles obtained for all the other time points 30 seconds apart for the duration of imaging are obtained and normalized using the mean intensity of the cell to

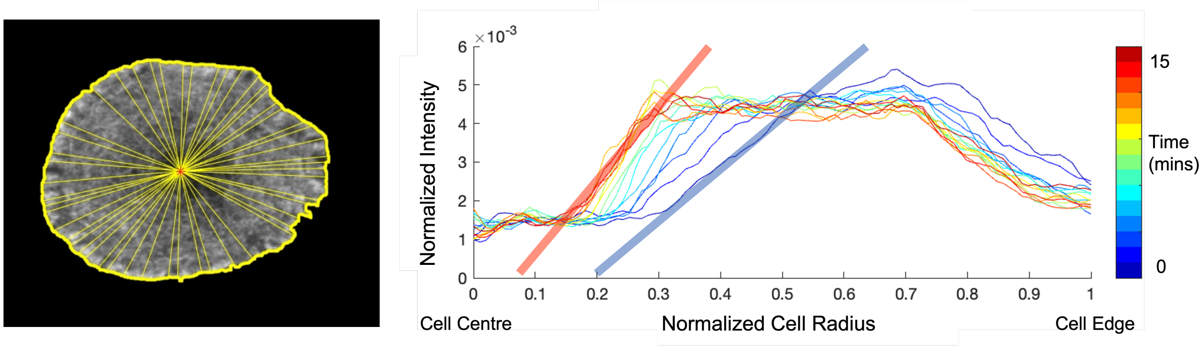


Figure SI-3: A example for calculating the slope of center to plateau F-actin distribution.

consider the effects of photobleaching. The resultant normalized line profile curves are now representative of how actin distribution changes over time inside the cell (Fig. SI-3, right). The intensity values typically increase from cell center shown by the centroid, which is the actin depletion region towards the cell edge before becoming coming stable at the maximum and then falling off. The steeply increasing region of the line profile curves are linearly fit (Fig. SI-3, right - shaded red and blue lines) to find their slope at each time point and the changing slope values over time are then compared among different chemical perturbations.
